# RNA localization to nuclear speckles follows splicing logic

**DOI:** 10.1101/2025.05.28.656493

**Authors:** Li Wen, Mauricio A. Arias, Xinqi Fan, Sneha Paul, Susan E. Liao, Marek Sobczyk, Oded Regev, Jingyi Fei

## Abstract

Nuclear speckles are membraneless organelles implicated in multiple RNA processing steps. In this work, we systematically characterize the sequence logic determining RNA localization to nuclear speckles. We find extensive similarities between the speckle localization code and the RNA splicing code, even for transcripts that do not undergo splicing. Specifically, speckle localization is enhanced by the presence of unspliced exon-like or intron-like sequence features. We demonstrate that interactions required for early splicesomal complex assembly contribute to speckle localization. We also show that speckle localization of isolated endogenous exons is reduced by disease-associated single nucleotide variants. Finally, we find that speckle localization strongly correlates with splicing kinetics of splicing-competent constructs and is tightly linked to the decision between exon inclusion and skipping. Together, these results suggest a model in which RNA speckle localization is associated with the formation of the early spliceosomal complex and enhances the efficiency of splicing reactions.

**Highlights:**

- Sequences containing hallmarks of pre-mRNA dictate speckle localization
- RNA speckle localization is coupled to early spliceosome assembly
- Disease-associated single nucleotide variants reduce localization of isolated exons
- RNA speckle localization strongly correlates with splicing kinetics

Graphical Abstract

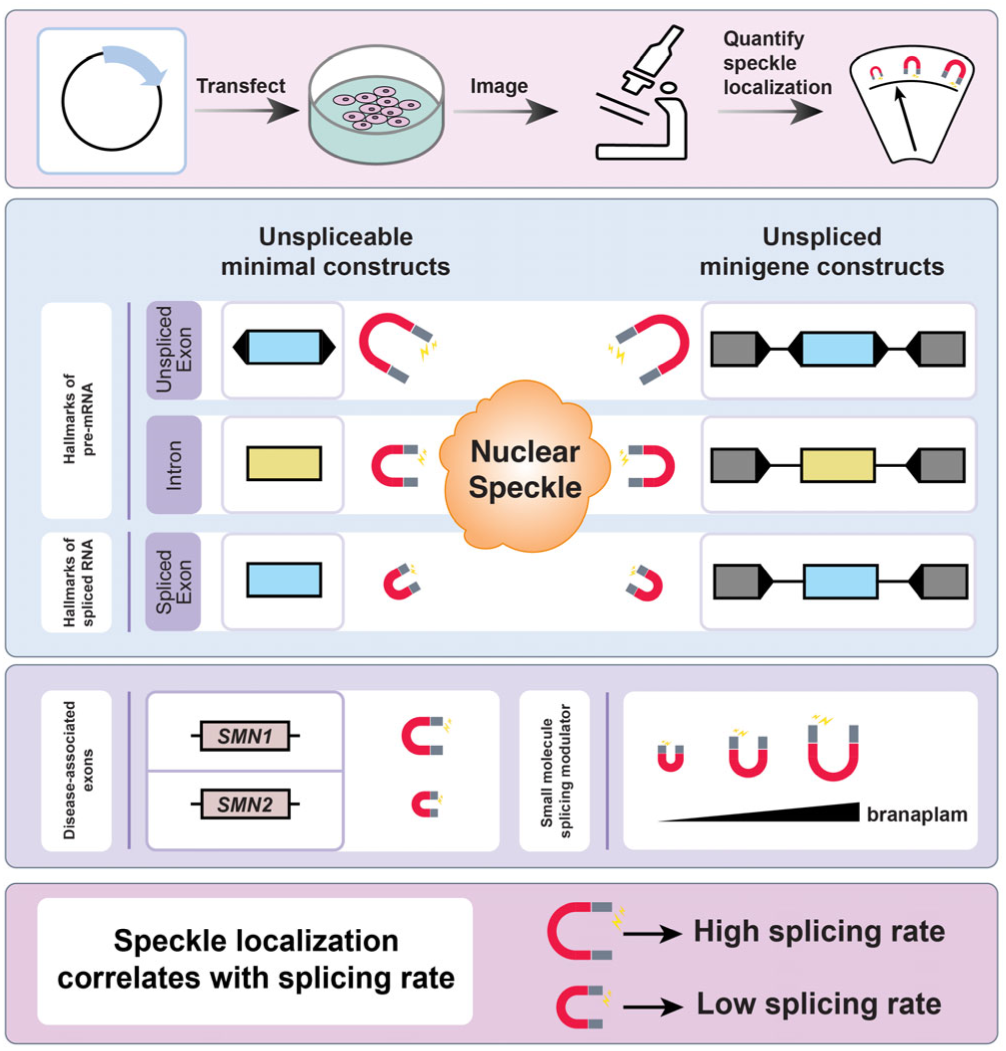

## Introduction

RNA localization to subcellular organelles is tightly linked to RNA processing. For instance, ribosomal RNA is processed and modified within the nucleolus and the biogenesis of small nuclear ribonucleoproteins (snRNPs) occurs in Cajal bodies.^1,2^ In these and other cases, identifying the RNA populations that localize to a particular organelle provides critical insights into the organelle’s functional roles.

A fraction of the nuclear transcriptome is localized to nuclear speckles, a prominent nuclear membraneless organelle.^3–6^ Human cells typically contain around 20-40 nuclear speckles ranging in size from a few hundred nanometers to a few microns. The nuclear speckle proteome contains components of the spliceosome and is enriched in splicing factors from the SR protein family.^6–8^ Consistent with its proteome composition, nuclear speckles have been implicated in several RNA processing steps, notably in facilitating splicing.^9–11^ In line with this potential functional role, speckle disruption leads to RNA splicing perturbation, including a global increase in intron retention and in exon skipping.^12,13^ However, the exact role of speckles in splicing remained unclear.

Recent studies of the nuclear speckle transcriptome by us and others identified multiple correlations between speckle localization and transcript properties.^12,14^ Speckle-enriched transcripts often have post-transcriptionally excised or retained introns, and typically contain short introns and high GC content. Speckle enrichment was also found to correlate with splice site strength, the presence of SR protein binding motifs in exons and introns, and with exon length.^12^ While these transcriptomic studies all point to correlations between splicing-related features and speckle localization, they do not establish any causality. Indeed, one challenge is that measurements of transcript speckle localization are likely affected not only by a transcript’s intrinsic affinity to speckles, but also by its splicing and nuclear export rate. A second challenge is that the position of a gene within the nucleus likely affects its transcripts’ speckle localization, as supported by a moderate correlation between gene distance to speckles and transcript speckle localization.^12,14^ Given that genes with high GC content and short introns are known to reside in the speckle-rich interior region of the nucleus,^15^ some of the correlations found in those transcriptomic studies are therefore likely due to the genome’s organization. In summary, decoupling these various contributions to transcript speckle localization is challenging with existing systems, limiting insights into speckle function.

Here, we systematically measure the speckle localization of synthetic reporter constructs. The use of a synthetic system with designed sequences allows us to probe transcripts that do not undergo splicing or export, which could otherwise complicate interpretation of the results. It also allows us to establish causal relationships between sequence and localization, as opposed to mere correlations. Using this approach, we characterize the “speckle localization code,” the sequence logic code determining nuclear speckle localization.

## Results

### SR motif-rich RNAs exhibit directional splice site-dependent nuclear speckle localization

To dissect the sequence determinants of transcript speckle localization, we started by generating a series of “minimal” reporter constructs that are not expected to undergo splicing (Figure 1A). Since SR proteins are enriched in nuclear speckles and were shown to affect speckle localization in previous reporter-based studies,^16,17^ we included a central region enriched in SR protein binding sites, as designed in our previous work.^8^ In more detail, we started with a sequence consisting of repeats of a characterized SRSF1 binding motif. We then randomly mutated 30% of the nucleotides, while ensuring that the resulting sequence is still enriched in SR binding motifs and, crucially, that no splice sites were generated.

**Figure 1.**
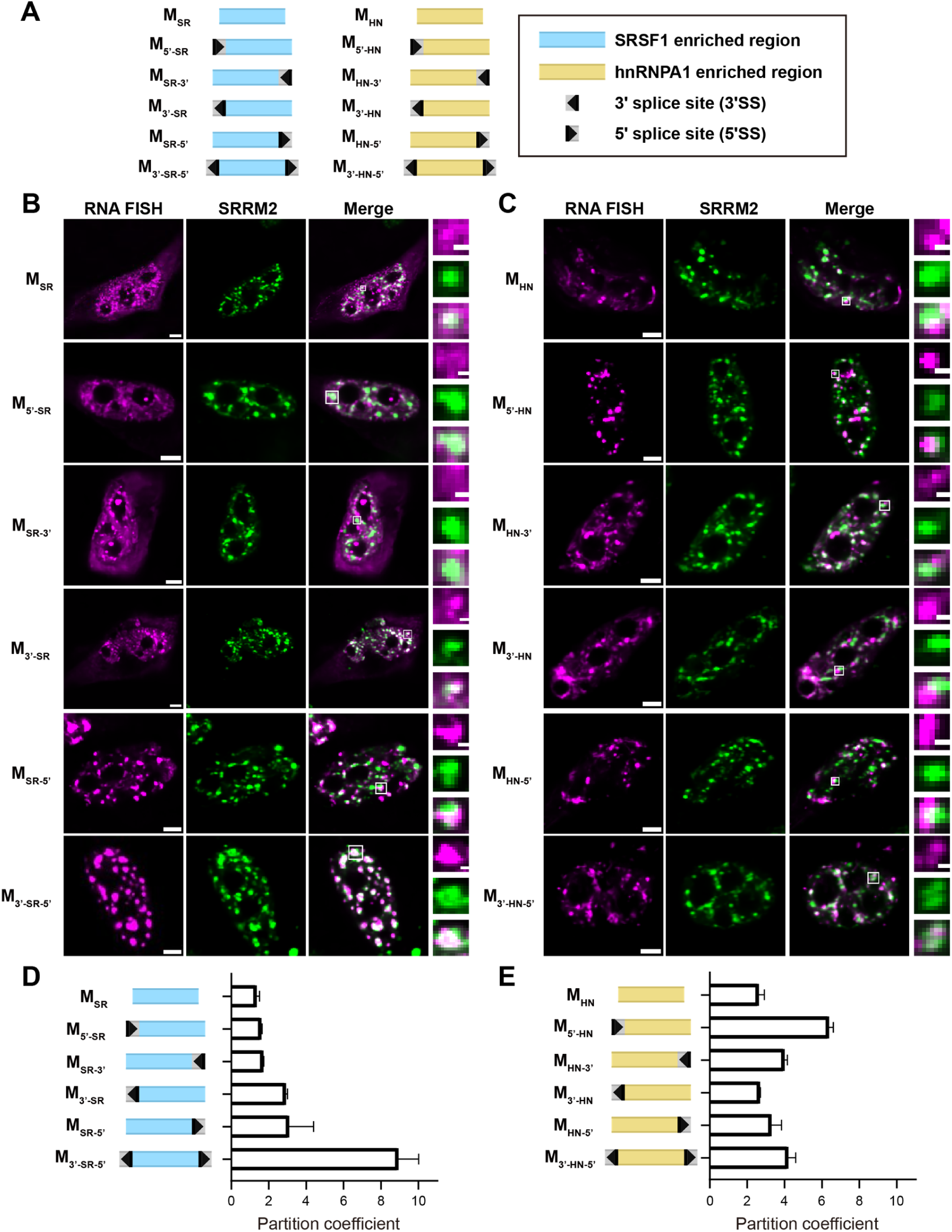
Localization of single-module unspliceable minimal constructs relative to nuclear speckles. (A) Schematic illustration of single-module unspliceable minimal constructs. (B–C) Representative RNA FISH image of minimal constructs containing SR motifs (B) and hnRNP motifs (C). RNA FISH signals (labeled with AF647) are shown in magenta. Immunofluorescence signals of SRRM2 (labeled with AF488) are shown in green. Scale bar (white line) represents 5 μm in images showing a single cell and 1 μm in zoomed-in images showing a single speckle. (D–E) Partition coefficients of minimal constructs containing SR motifs (D) and hnRNP motifs (E). Error bars report the standard deviation from 2 biological replicates, with each replicate including 69–205 cells.

We next generated six reporters by adding various combinations of 3’ and 5’ splice site sequences around this central region. We excluded the combination of an upstream 5’ splice site with a downstream 3’ splice site as such a construct might get spliced. The constructs, which are all driven by a tet-responsive promoter, were next transfected into a HeLa cell line with stably expressed Tet-regulated transactivator Tet-On 3G, and RNA expression was induced with doxycycline for 2 h. To rule out any unexpected products from our constructs (due to, e.g., cryptic transcription, cryptic splicing, or cryptic polyadenylation), we performed transcriptome-wide long-read polyA RNA sequencing, capturing entire transcripts from their 5’ end to their polyA tail (Figure S1). For almost all constructs, the vast majority of transcripts are precisely as intended. In a few constructs, however, we found a more significant fraction of unintended transcripts. For instance, a small fraction of transcripts were generated from a previously characterized cryptic transcription site inside the bacterial origin of replication.^18^ In a few other cases, some readthrough transcripts extended beyond the polyA signal and were spliced into downstream sequences, typically in constructs with unmatched 5’ splice sites, which are known to suppress nearby polyadenylation sites.^19^ We verified that these unintended transcripts do not affect our measurements or conclusions (Figure S2, Supplemental notes).

We then measured the speckle localization of these constructs. We labeled RNA transcripts via fluorescence in situ hybridization (FISH) using fluorophore-labeled DNA oligos, and immunostained nuclear speckles using antibodies against the marker protein SRRM2 (Figure 1B). Minimal background FISH signal was observed from cells that were not successfully transfected, confirming the specificity of the FISH probes (Figure S2A). To quantitatively report the speckle localization propensity of each transcript, we computed a “partition coefficient” (P), defined as the ratio of RNA FISH intensity within speckles to that in the nucleoplasm. The partition coefficient therefore reflects the RNA concentration within speckles relative to that in the nucleoplasm. The constructs exhibit a wide range of partition coefficients (Figure 1D). These partition coefficients were not correlated with the constructs’ expression levels (Figure S3A). In addition, within each construct, the partition coefficient is not correlated with RNA expression level at the single-cell level (Figure S3B). Together, this establishes the partition coefficient as an intrinsic property of each construct rather than being driven by transcript abundance.

We found that speckle localization of these SR motif-enriched transcripts depends on the presence of splice sites as well as their relative position (Figure 1D). Specifically, despite the enrichment of SR proteins in speckles, the SR motif-rich region by itself shows low speckle localization. Moreover, introducing a 5’ splice site upstream or a 3’ splice site downstream of the SR motif-rich region did not increase speckle localization. In contrast, introducing a 3’ splice site upstream or a 5’ splice site downstream of the SR motif-rich region noticeably increased speckle localization. Finally, including both splice sites at those positions significantly boosted speckle localization. Together, these results indicate that speckle localization is specifically promoted by a defined positioning of splice sites relative to SR motif-rich regions.

These positional behaviors are reminiscent of the sequence features that characterize unspliced exons.^20^ These features include an upstream 3’ splice site and a downstream 5’ splice site, and often an enrichment of SR protein binding motifs, which serve as exonic splicing enhancers.^20^ Our results show that RNA with such a sequence combination (M_3’-SR-5’_) shows a high speckle localization propensity. When this sequence combination is only partly formed (as in M_3’-SR_ and M_SR-5’_), the RNA shows an intermediate speckle localization propensity. In contrast, when SR motifs are not flanked by splice sites (as in M_SR_), the RNA shows the lowest speckle localization. We hypothesize that this low speckle enrichment is due to the SR motifs in this sequence context being identified as spliced, mature mRNAs (as those mRNAs typically contain SR-enriched exonic sequences but no splice sites). Indeed, M_SR_ showed a high cytoplasmic fraction, consistent with it being exported as mature mRNA (Figure S4). In addition, SR motifs are known to inhibit upstream 5’ splice sites and downstream 3’ splice sites, thereby acting as intronic silencers;^20–22^ this explains why M_5’-SR_ and M_SR-3’_ have a low speckle localization propensity, similarly to M_SR_. Consistently, M_SR-3’_ also showed a high cytoplasmic fraction (Figure S4). The low cytoplasmic fraction of M_5’-SR_ (Figure S4) is likely due to the observation that an unpaired 5’ splice site inhibits RNA export.^23^ In summary, these results indicate that sequence features of pre-mRNA promote speckle localization, whereas sequence features of mature spliced mRNA reduce speckle localization.

### hnRNP motif-rich RNAs exhibit inherent speckle localization propensity

Contrary to SR proteins, hnRNP proteins are generally not enriched in nuclear speckles and even slightly depleted.^8^ These two families of splicing regulators typically have opposing effects on mRNA splicing, as suggested by the enrichment of SR binding motifs in exons and the enrichment of hnRNP binding motifs in introns.^20^ In addition, while exon-bound SR proteins demonstrate splicing enhancer activity, promoting exon inclusion, exon-bound hnRNP proteins act as splicing silencers, promoting exon skipping by effectively identifying that region as an intron.^20^ We therefore next generated an analogous set of six constructs in which we replaced the SR motif-rich region with an hnRNP motif-rich region of the same length (Figure 1A). This region was designed as before, but starting with a sequence containing hnRNPA1 motifs.^8^ We again used long-read sequencing to rule out the presence of unwanted cryptic transcripts (Figures S1 and S2, Supplemental notes).

In contrast to the SR motif-rich constructs, transcripts from the hnRNP motif-rich constructs exhibited intermediate levels of nuclear speckle localization that are mostly independent of splice sites (Figure 1C and E). All hnRNP constructs, including the one without any splice sites (M_HN_), exhibit a partition coefficient that is higher than that of the SR construct M_SR_. Moreover, inspecting the microscopy images revealed that this intermediate partition coefficient results from localization to the speckle periphery (Figure 1C), consistent with a previous observation via super-resolution FISH imaging,^8^ and possibly related to a depletion of hnRNP proteins inside speckles.^8,24,25^

We note that M_5’-HN_ transcripts exhibit higher speckle localization than the other hnRNP motif-containing constructs and are significantly enriched in the interior of speckles rather than on their periphery (Figure 1C and E). To ensure that this higher partition coefficient is not due to cryptic sequence elements, we performed two validations. First, we removed a polypyrimidine tract-like sequence and a 3’ splice site-like sequence which we identified as part of the constant region at the 5’ end of the transcript. In both cases, the partition coefficient remained similar (Figure S5). Second, we ruled out activation of a cryptic 3’ splice site downstream of the polyA signal by linearizing the M_5’-HN_ plasmid, again observing no change in partition coefficient (Figure S2B and F, Supplemental notes). These experiments indicate that the higher partition coefficient of M_5’-HN_ is not due to activation of cryptic splice sites; instead, it is the combination of a 5’ splice site followed by hnRNP motifs (as in M_5’-HN_) that promotes speckle localization. We speculate that this might be due to the region upstream of the 5’ splice site, aided by the downstream intron-like hnRNP region, being identified as a first exon.

In summary, these results indicate that hnRNP motifs provide transcripts with an inherent speckle affinity. Contrary to SR motifs, which depend on splice sites for speckle localization, hnRNP motifs showed speckle localization even in the absence of splice sites. These results suggest that hnRNP motifs, which are enriched in introns,^20^ can contribute to intron recognition via speckle localization.

Collectively, these results on SR motif-enriched and hnRNP motif-enriched constructs suggest that RNA speckle localization is driven by the directional combination of splice sites and binding sites of splicing regulators. In all cases considered, sequences containing hallmarks of pre-mRNA (specifically, an unspliced exon or an intronic-like sequence) tend to drive speckle localization. Sequences resembling mature spliced mRNA (which does not contain unspliced exons nor intronic-like sequences) are not enriched in nuclear speckles.

### Interactions with U1 snRNP and U2AF contribute to RNA speckle localization

While our data suggest that RNA localization to speckles does not require a splicing reaction, its dependence on the presence of splicing-related sequence features is indicative of a synergy with splicing regulators and spliceosomal components. To test whether recognition of the 5’ splice site by the spliceosomal U1 snRNP complex impacts speckle localization, we first mutated the strong 5’ splice site in M_3’-SR-5’_ to a weak splice site, as quantified by the MaxEnt score^26^ (Figure 2A). Weakening the 5’ splice site reduced speckle localization (Figure 2B). Importantly, co-transfection with a mutant U1 snRNA complementary to the weak 5’ splice site, but not with wild type U1 snRNA, rescued speckle localization of the weak 5’ splice site variant (Figure 2C).

**Figure 2.**
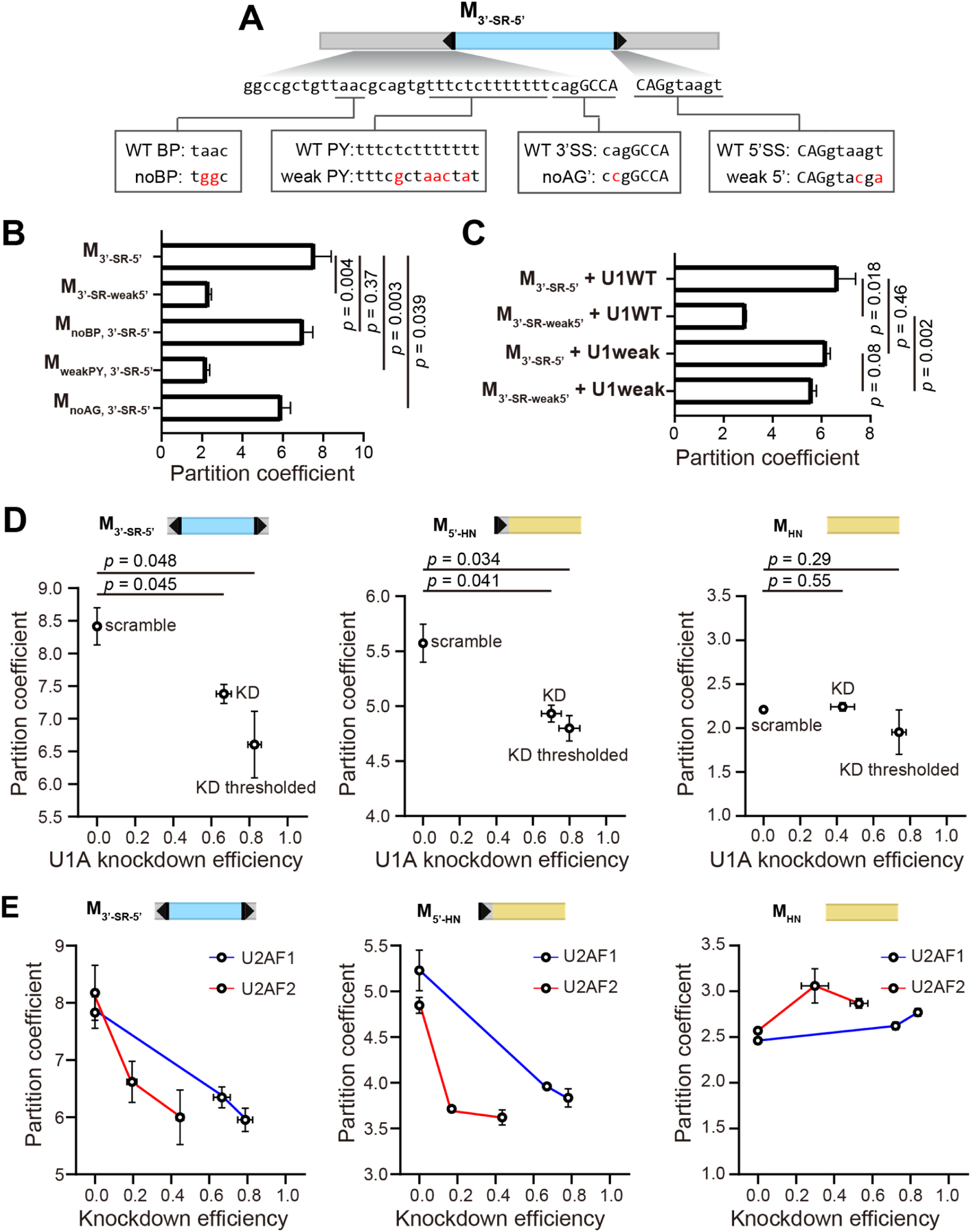
Interactions with U1 snRNP and U2AF contribute to speckle localization of minimal constructs. (A) Schematic illustration of sequence perturbations introduced to M_3’-SR-5’_. BP and PY indicate the branch point and polypyrimidine tract. (B) Partition coefficients of various M_3’-SR-5’_ mutant constructs in comparison with WT. (C) Partition coefficients of WT M_3’-SR-5’_ and M_3’-SR-weak5’_ in the presence of plasmid-encoded WT U1 snRNA or mutant U1 snRNA with sequence complementary to the 5’ splice site of M_3’-SR-weak5’_. (D) Scatter plot of partition coefficient vs. U1A knockdown efficiency for M_3’-SR-5’_, M_5’-HN_, and M_HN_ in the presence of scramble siRNA or siRNA targeting the *U1A* gene. “KD” marks the value averaged from all cells in the U1A siRNA treated sample. “KD thresholded” marks the value averaged from the subset of cells with U1A knockdown efficiency over 70%. (E) Scatter plot of partition coefficient vs. knockdown efficiency for M_3’-SR-5’_, M_5’-HN_, and M_HN_ in the presence of scramble siRNA or siRNA targeting the *U2AF1* (blue line) or *U2AF2* (red line) genes. Data point corresponding to 0% knockdown efficiency represents the value averaged from all the cells in the scramble siRNA-treated sample. The two data points corresponding to >0% knockdown efficiency represent values averaged from the subsets of cells with bottom 50% and top 50% knockdown efficiency respectively. Error bars report the standard deviation from 2-4 biological replicates. In (B) and (C), 29-300 cells per replicate were included. In (D) and (E), when no threshold was applied, each replicate included measurements from 69-686 cells; with thresholding, 15-312 cells per replicate were included. *P*-values were calculated from unpaired two-tailed t-test.

We further examined two constructs with high speckle localization, M_3’-SR-5’_ and M_5’-HN_, under siRNA knockdown of the U1 snRNP component U1A. We verified that U1A knockdown did not have a significant impact on speckle morphology (Figure S6). Since knockdown efficiency of the abundant U1A protein showed heterogeneity across the cell population, we co-stained U1A together with SRRM2 and RNA FISH, and quantified the speckle partition coefficient among high knockdown efficiency cells (Figure 2D). Our results demonstrate a reduction of partition coefficient as knockdown efficiency increases for both M_3’-SR-5’_ and M_5’-HN_, but not for M_HN_, which lacks a 5’ splice site. These results suggest that transcript localization to speckles is modulated by the efficiency of 5’ splice site recognition by U1 snRNP.

To test whether recognition of the 3’ splice site also impacts RNA speckle localization, we introduced three perturbations to M_3’-SR-5’_: mutating the branch point, weakening the polypyrimidine tract, and mutating the conserved AG dinucleotide at the 3’ splice site (Figure 2A). We observed a large reduction in partition coefficient upon weakening the polypyrimidine tract, and a smaller one with the AG dinucleotide mutation. In contrast, the branch point mutation did not show a significant impact (Figure 2B), even though we verified that no other strong predicted branch point remains.^27^ These results indicate that transcript localization is associated with the recognition of the polypyrimidine tract and 3’ splice site, but not with the recognition of the branch point.

The polypyrimidine tract and the 3’ splice site AG are initially recognized by U2AF2 and U2AF1 respectively. We therefore further examined the speckle localization of M_3’-SR-5’_ and M_5’-HN_ upon knocking down U2AF1 or U2AF2 (Figure 2E). The partition coefficient of M_3’-SR-5’_ was especially sensitive to U2AF2 knockdown, showing a large decrease even for relatively mild knockdown efficiencies. In contrast, the partition coefficient of M_HN_, which does not have a polypyrimidine tract or a 3’ splice site sequence, is not affected by U2AF2 and U2AF1 knockdown. This observation is consistent with the above result that perturbation of the polypyrimidine tract has a larger impact on speckle localization. We note that M_5’-HN_ transcripts also demonstrated a reduction in partition coefficient upon U2AF1 and U2AF2 knockdown even though they lack polypyrimidine tracts and 3’ splice sites; the reasons for that are not clear.

Together, these results suggest that transcript speckle localization may occur during early spliceosome assembly, more specifically during the formation of the spliceosomal E complex. Indeed, 5’ splice sites are recognized by U1 snRNP during early spliceosomal assembly, consistent with their strong effect on localization observed here. Moreover, 3’ splice site mutations are known to block spliceosomes at a late catalytic step,^28^ consistent with their observed weak effect on partition coefficient. This hypothesis is also consistent with our recent work, showing that splicing inhibition using Pladienolide B (Plad B), which stalls spliceosomes at A complex,^29^ does not significantly affect speckle localization of pre-mRNA transcripts (Figure S2E).^12^

### Speckle localization depends non-additively on exonic sequence and is affected by interaction with SRSF1 and hnRNPA1

Our results show that M_3’-HN-5’_ has an intermediate partition coefficient whereas M_3’-SR-5’_ has a high one. To further investigate how the balance between SR and hnRNP binding to the exonic sequence affects speckle localization, we generated two “trajectory” series (Figure 3A and C). These series interpolate between M_3’-HN-5’_ and M_3’-SR-5’_ by successively replacing hnRNP-enriched regions with SR-enriched ones. As expected, we observed an increase in nuclear speckle partition coefficient along both series. Interestingly, we noticed a somewhat abrupt transition between M_A3_ and M_A4_, and between M_B3_ and M_B4_ (Figure 3B and D), showing that the effect of RBP binding motifs on speckle localization is not additive, and mirroring observations on splicing decisions.^30^

**Figure 3.**
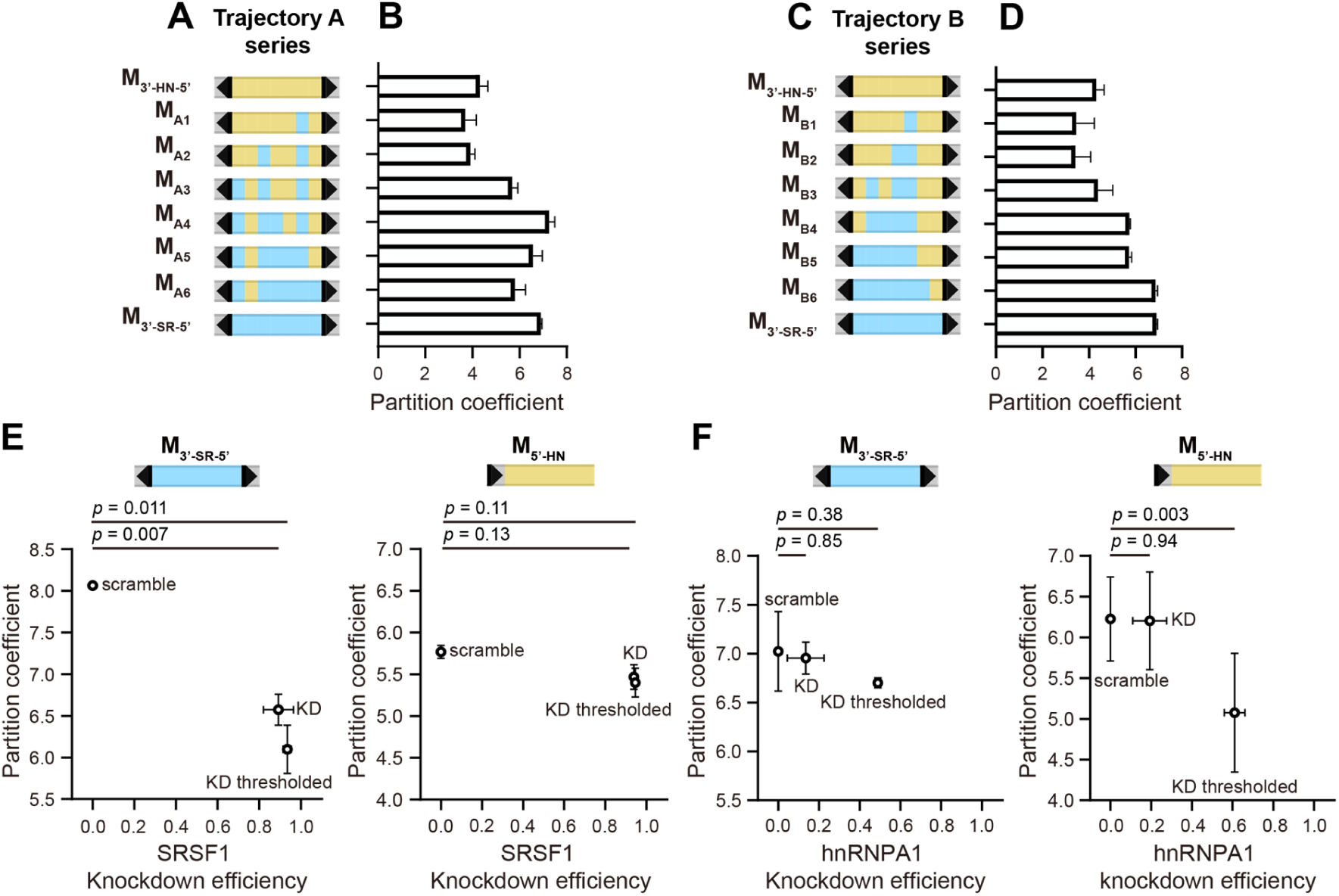
Speckle localization is affected by interaction with SRSF1 and hnRNPA1. (A), (C) Schematic illustration of trajectory series A (A) and B (C). Color gradient from yellow (most enriched in hnRNP motifs in M_3’-HN-5’_) to cyan (most enriched in SR motifs in M_3’-SR-5’_) represents a gradual increase in the fraction of SR motifs in the sequence. (B), (D) Partition coefficients of trajectory series A (B) and B (D). Data for F_3’-SR-5’_ and F_3’-HN-5’_ are the same in both plots. (E–F) Scatter plot of partition coefficient vs. SRSF1 (E) or hnRNPA1 (F) knockdown efficiency for M_3’-SR-5’_ and M_5’-HN_ in the presence of scramble siRNA or siRNA targeting *SRSF1* or *hnRNPA1* gene. “KD” marks the value averaged from all cells in the SRSF1 or hnRNPA1 siRNA treated sample. “KD thresholded” marks the value averaged over the subset of cells with knockdown efficiency over 90% for SRSF1 and 50% for hnRNPA1. Error bars report the standard deviation from 2-8 biological replicates. In (B) and (D), 34-840 cells per replicate were included. In (E) and (F), when no threshold was applied, each replicate included measurements from 78-651 cells; with thresholding, 18-444 cells per replicate were included. *P*-values were calculated from unpaired two-tailed t-test.

To further investigate the contribution of SRSF1 and hnRNPA1 to RNA localization, we imaged M_3’-SR-5’_ and M_5’-HN_ transcripts upon SRSF1 and hnRNPA1 KD. As expected, SRSF1 KD reduced the partition coefficient of M_3’-SR-5’_ transcripts, but had insignificant impact on M_5’-HN_ (Figure 3E). Similarly, hnRNPA1 KD reduced the partition coefficient of M_5’-HN_, but had insignificant impact on M_3’-SR-5’_. We note that the impact of hnRNPA1 KD was milder than that of SRSF1 KD, likely due to the lower KD efficiency achieved in our experimental conditions (Figure 3F).

### The SRSF1 motif dominates the hnRNPA1 motif for nuclear speckle localization

We next considered cases where SR and hnRNP motifs are both present in the RNA. We generated a series of minimal “dual-module” constructs (Figure 4A), using different combinations of splice sites, SRSF1, and hnRNPA1 motifs. We again used long-read sequencing to rule out cryptic transcripts that can affect our results (Figure S1). Transcripts from the dual-module minimal constructs displayed a wide range of partition coefficients (Figure 4A). We then compared the partition coefficients of the dual-module constructs to those of the single-module constructs containing the analogous SR/hnRNP motifs and splice sites combination and orientation. Interestingly, we found that the partition coefficients of the dual-module transcripts are strongly correlated with the analogous SR motif-containing single-module transcripts (Figure 4B), but are weakly correlated with the hnRNP motif-containing single-module transcripts (Figure 4C). These comparisons suggest that when both motifs are present, SR motif has a dominant effect on speckle localization.

**Figure 4.**
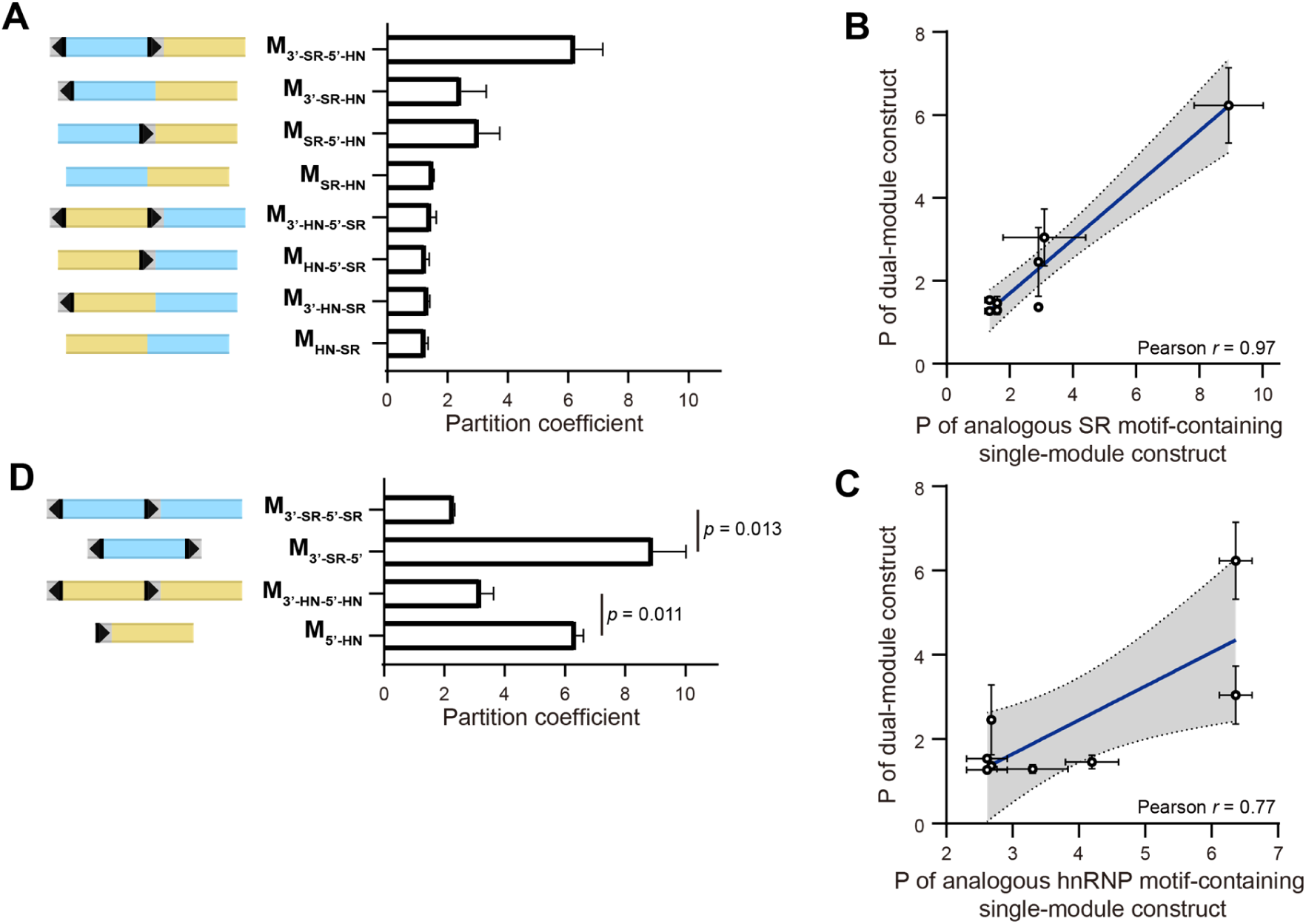
Comparison between single-module and double-module minimal constructs. (A) Schematic illustration of dual-module minimal constructs containing both SR and hnRNP motifs and corresponding partition coefficients. (B) Comparison between partition coefficients of dual-module constructs in (A) and corresponding single-module constructs containing SR motifs in Figure 1D. (C) Comparison between partition coefficients of dual-module constructs in (A) and corresponding single-module constructs containing hnRNP motifs in Figure 1E. (D) Comparison between partition coefficients of dual-module constructs containing two regions enriched in SR or hnRNP motifs and corresponding single-module constructs. Error bars report the standard deviation from 2-3 biological replicates, with each replicate including 36–188 cells. *P*-values were calculated from unpaired two-tailed t-test. Pearson correlation coefficients (*r*) are shown in (B) and (C).

We additionally generated a construct containing two SR motifs (M_3’-SR-5’-SR_) and another containing two hnRNP motifs (M_3’-HN-5’-HN_) (Figure 4D). M_3’-SR-5’-SR_ exhibited a significantly lower partition coefficient than that of M_3’-SR-5’_ (Figure 4D). This result suggests that the downstream SR motif inhibits the recognition of the 5’ splice site, lowering the partition coefficient similarly to the disruption of 5’ splice site recognition (Figure 2B and D). The second construct, M_3’-HN-5’-HN_, exhibited an intermediate partition coefficient significantly lower than that of M_5’-HN_. This result similarly suggests that the upstream hnRNP motifs inhibit the recognition of the 5’ splice site (Figure 4D). Together, these results indicate that the efficient recognition of 5’ splice sites benefits from an upstream sequence enriched in SR motifs and a downstream sequence enriched in hnRNP motifs. In the absence of such a sequence combination, the 5’ splice site recognition is compromised. The position-dependent effects of SR and hnRNP motifs on transcript speckle localization is in line with their known position-dependent effects as splicing enhancers and suppressors.^31^

### Context-dependent nuclear speckle localization of minimal constructs is largely preserved in pre-mRNA from minigene constructs

We next examined whether the context-dependent speckle localization observed in RNA from the minimal constructs also exists in pre-mRNA from splicing-competent constructs. We therefore created a series of minigene constructs by inserting the minimal constructs, including the two trajectory series, into the intron of a two-exon minigene (Figures 5A, 5B for Group I and S7A for Groups II-III). We first analyzed the splicing outcome of these minigenes, allowing us to identify what part of each minigene gets identified as an intron and removed during the splicing process (Figures 5C and S7B), which we can image to report the localization of pre-mRNA. In addition, to avoid confounding signal contributed by the spliced intron lariats, we computed the abundance ratio of the intron-exon junction to that of an intron-internal region using RT-qPCR (Table S1). This ratio was never smaller than 70%, suggesting that all introns are efficiently degraded. We designed probes against the examined intronic regions of each construct, allowing us to specifically investigate the localization of unspliced pre-mRNA. We quantified the pre-mRNA partition coefficient (Figures 5D and S7C) and compared it with the partition coefficient of the corresponding minimal construct (Figure 5E, Table S2). Interestingly, the comparison indicated that despite the additional sequence context provided by the upstream and downstream introns and exons, the nuclear speckle localization propensity of the minimal constructs can largely predict that of the minigene pre-mRNAs (Pearson’s *r* = 0.87).

**Figure 5.**
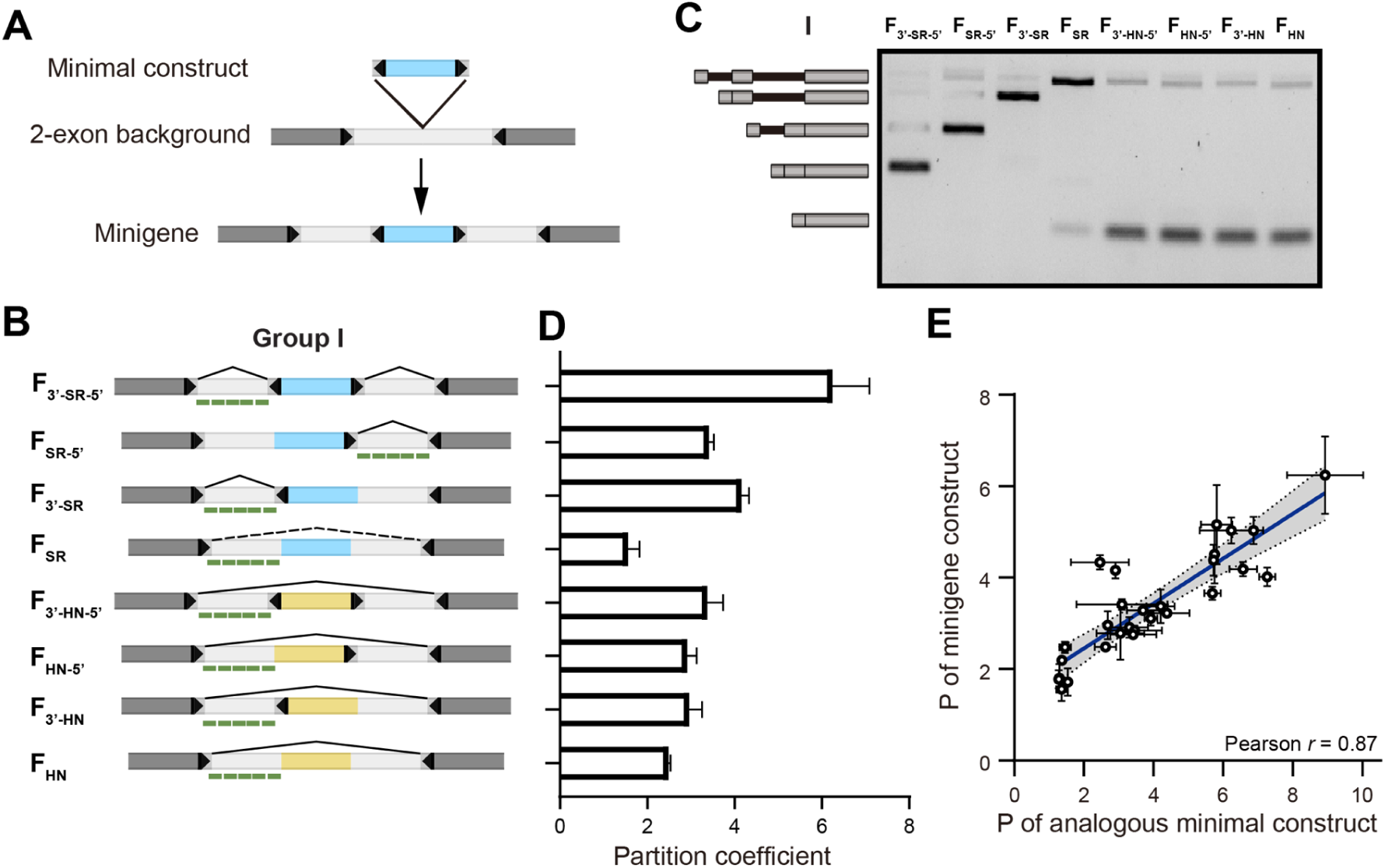
Localization of pre-mRNA from minigene constructs. (A) Schematic illustration of generation of minigene constructs from minimal constructs. (B) Schematic illustration of minigene constructs containing single-module minimal construct sequences (Group I) with splicing outcome marked using solid lines (presenting major splicing products) or dashed lines (representing partial splicing products). The other three groups (II, III, and IV) are shown in Figure S7A. Green bars mark the FISH probe targeting sites in each construct. (C) RT-PCR gel electrophoresis analysis of splicing products for minigene constructs in (B). Analyses for the other three groups (II, III, and IV) are shown in Figure S7B. (D) Partition coefficients of pre-mRNAs from minigene constructs in (B). Analyses for the other three groups (II, III, and IV) are shown in Figure S7C. (E) Comparison between partition coefficients of minigene pre-mRNAs and corresponding minimal constructs. Error bars report the standard deviation from 2-3 biological replicates, with each replicate including 23–204 cells. Pearson correlation coefficient (*r*) is shown in (E).

The speckle localization propensity of these minigene pre-mRNAs and their splicing outcomes elucidate and provide further support to our hypothesis that the presence of unspliced exons or intron-like sequences promotes speckle localization. For instance, similarly to RNA from the minimal construct M_3’-SR-5’_, pre-mRNA from F_3’-SR-5’_ shows a high speckle localization propensity (Figure 5B and D). As expected given the role of SR proteins as exonic enhancers, when this minigene is spliced, the SR motif-containing region is identified as an exon and is included in the mature mRNA (Figure 5C). This observation supports our hypothesis that the high speckle localization of both M_3’-SR-5’_ and F_3’-SR-5’_ is driven by the variable region being identified as an unspliced exon, even though no splicing takes place with the former. As another example, the four hnRNP-only minigenes (F_3’-HN-5’_, F_HN-5’_, F_3’-HN_, and F_HN_) show the same intermediate speckle localization propensity as the corresponding minimal constructs (Figure 5B and D). Moreover, as expected given the role of hnRNP proteins as exonic silencers, in all four minigenes, the hnRNP motifs are identified as part of the intron, leading to skipping of the entire region between the terminal exons (Figure 5C). This supports our hypothesis that the intermediate speckle localization of these four minimal constructs and four minigenes is driven by the hnRNP motif-containing region being identified as an intronic sequence. A final interesting example is F_SR_, whose pre-mRNA shows a poor speckle localization as was the case for M_SR_. Here, the RT-PCR assay confirmed that F_SR_ is largely unspliced (Figure 5C), consistent with intron-positioned SR motifs serving as intronic silencers and associated with intron retention. This observation suggests that the SR motifs interfere with the recognition of the middle intron,^22^ driving the low speckle localization. We inspected the remaining minigene constructs, as well as four additional minigene constructs containing a neutral sequence (N) not enriched in SR or hnRNP motifs (Figure S7A, Group IV). All cases agree with the general pattern that unspliced exons and intron-like sequences promote speckle localization.

### Speckle localization correlates with exon inclusion

Exon skipping is one of the most common forms of alternative splicing in the human transcriptome.^32^ We therefore used the trajectory minigene series to measure how the transition from exon skipping to exon inclusion is reflected in terms of speckle localization. Using RT-qPCR, we quantified the percentage spliced in (PSI) of those constructs (Figure 6A, Table S3). As previously mentioned, the first minigene in both series (F_3’-HN-5’_) shows excision of the entire middle region, i.e., exon skipping. The last minigene (F_3’-SR-5’_) shows strong exon inclusion. Along the trajectory, we observed a sharp transition in PSI starting around F_A2_ and F_B2_, consistent with the known switch-like behavior of splicing decisions. Beyond that transition, PSI saturated at close to 100% while the partition coefficient kept increasing, consistent with our above observation that knocking down SRSF1 protein reduces speckle localization of M_3’-SR-5’_. Interestingly, the increase in PSI observed at the transition point (F_A2_ and F_B2_) is not yet accompanied by a significant increase in partition coefficient (Figure 6A), indicating that PSI is more sensitive to exon sequence than speckle partition coefficient. Finally, although the general trend was of increasing speckle localization along the trajectory, we did notice a slight decrease with the first few minigenes of each series, a pattern also visible in the analogous minimal series (Figures 3A-D, S7C III). While the decrease might not be statistically significant, this behavior agrees with our earlier observation that the presence of SR motifs inside a recognized intron region (as in F_SR_) weakens its definition and reduces speckle localization. Once a critical threshold of SR motifs is reached, the region is identified as an exon, and additional SR motifs increase speckle localization and PSI. In summary, these results show that speckle localization is tightly linked to exon inclusion decisions.

**Figure 6.**
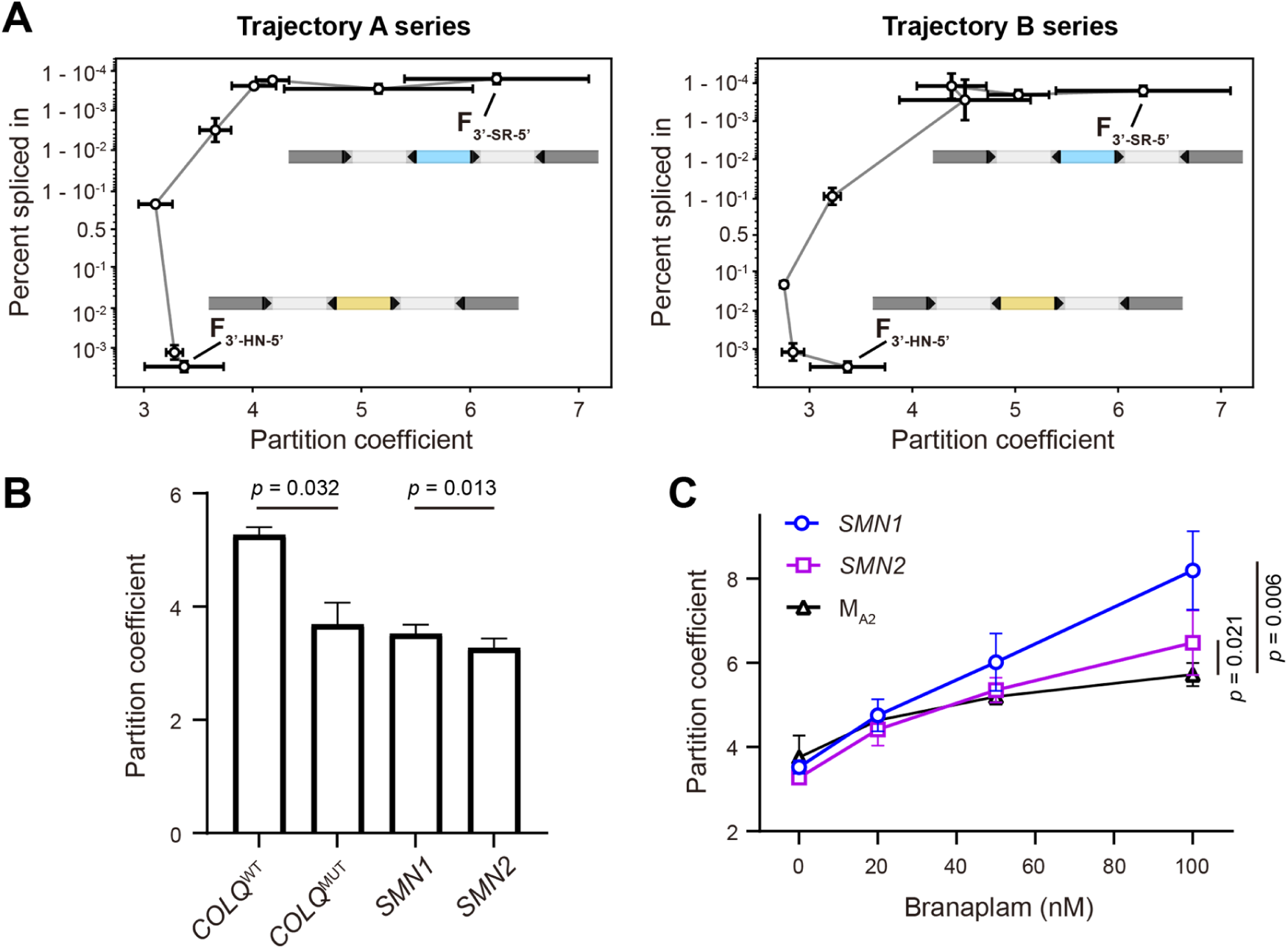
Speckle localization correlates with exon inclusion. (A) Correlation between partition coefficients of pre-mRNAs from minigene constructs for two trajectory series and the middle exon’s percent spliced in. Data for F_3’-HN-5’_ and F_3’-SR-5’_ are the same in both plots. The y-axis is shown on a logit scale. (B) Partition coefficients of minimal constructs containing WT (*COLQ*^WT^) and mutant (*COLQ*^MUT^) *COLQ* gene exon 16, as well as *SMN1* and *SMN2* gene exon 7. (C) Changes in the partition coefficients of *SMN1*, *SMN2,* and M_A2_ minimal constructs in the presence of branaplam. In (A), error bars in percentage spliced in report the standard deviation from 2 biological replicates, which are smaller than the symbol size. Error bars in partition coefficient report the standard deviation from 2 biological replicates, with each replicate including 23–88 cells. In (B) and (C), error bars report the standard deviation from 5-7 biological replicates, with each replicate including 114–1123 cells. *P*-values in (B) were calculated using unpaired two-tailed t-test. *P*-values in (C) were calculated using ANCOVA between *SMN1* and M_A2_, and between *SMN2* and M_A2_.

### Variants known to affect splicing also affect speckle localization of isolated endogenous exons

While our results highlight a close connection between speckle localization and splicing decisions, they are all based on designed sequences. We therefore asked whether endogenous sequences exhibit a similar speckle localization pattern. Specifically, mutations that perturb SR or hnRNP protein binding sites are often found in splicing-related genetic disorders, such as in the *SMN1/2* genes in spinal muscular atrophy and the *COLQ* gene in endplate acetylcholinesterase deficiency.

We therefore first considered the well-characterized exon 7 of *SMN1*.^33,34^ The human genome contains a nearly identical gene, *SMN2*. A single nucleotide difference at position +6 of exon 7 is enough to cause inefficient splicing of that exon in *SMN2* compared to *SMN1*, due to the loss of an SRSF1 binding site.^33^ We generated two minimal constructs expressing these short 54 nt exons with their flanking splice sites. Using nanopore long-read sequencing, we verified that the transcripts were expressed largely as expected without any splicing occurring, and that the minor fraction of unintended transcripts did not affect our imaging measurements (Figures S1, S2B, and S2G, Supplemental notes). Even though the two constructs differ in only one exon-internal nucleotide, imaging revealed a lower partition coefficient in *SMN2* exon 7 compared to *SMN1* exon 7 (Figure 6B). This result is consistent with the known splicing behavior of these exons in their endogenous context.

To further demonstrate the correlation between splicing and speckle localization, we repeated the imaging experiments after treating cells with increasing concentrations of branaplam, a small molecule developed to increase exon 7 inclusion.^35^ The molecule enhances the binding of U1 snRNP to the 5’ splice site of exon 7, and is reported to be highly specific.^35,36^ We found that branaplam treatment increased speckle localization of both *SMN1* exon 7 and *SMN2* exon 7 in a dose-dependent manner, as they have the same 5’ splice site sequence. As a control, we used M_A2_, which has a different 5’ splice site sequence and a similar partition coefficient, and found a smaller increase in partition coefficient upon treatment (Figure 6C). This observation is consistent with the 5’ spice site sequence dependence of branaplam.^36^

We next performed a similar experiment with the *COLQ* gene. A single nucleotide mutation inside exon 16 leads to exon skipping by abolishing an SRSF1 binding site while generating a novel hnRNPH binding site.^37^ We again generated two minimal constructs expressing the isolated exons with some of their flanking genomic sequences. We verified that no splicing takes place in these constructs, and imaged them as before. We found that the mutant construct exhibits a much lower speckle localization compared to the wild type (Figure 6B), in agreement with the mutation’s effect on splicing.^37^

Together, these data show that results obtained with synthetic sequences also apply to endogenous sequences. They also highlight how subtle features of the splicing code (namely, the effect of single nucleotide mutations that do not form or abolish splice sites) are recapitulated by speckle localization, even though no splicing is taking place.

### Nuclear speckle localization correlates with nuclear RNA degradation and splicing kinetics

Given the observed pre-mRNA context-dependent nuclear speckle localization driven by the splicing regulators, we wondered whether the propensity of speckle localization impacts splicing rate. We set up a platform combining imaging-based kinetic measurement with computational modeling to determine RNA processing parameters: transcription rate, splicing rate, and degradation rate (of pre-mRNA and spliced mRNA) (Figure 7A and B). We randomly selected a subset of minigene constructs exhibiting a wide range of speckle partition coefficients. Two-color FISH probes were used to target the intron and exon regions of the transcript (Figure 7A). The exonic probe signal integrated over the entire cell reports the sum of pre-mRNA and spliced mRNA abundance for each construct, whereas the intronic probe signal reports the total abundance of pre-mRNA and (undegraded) spliced intron lariat. Transcription was induced at time zero and FISH signals were recorded as a function of time.

**Figure 7.**
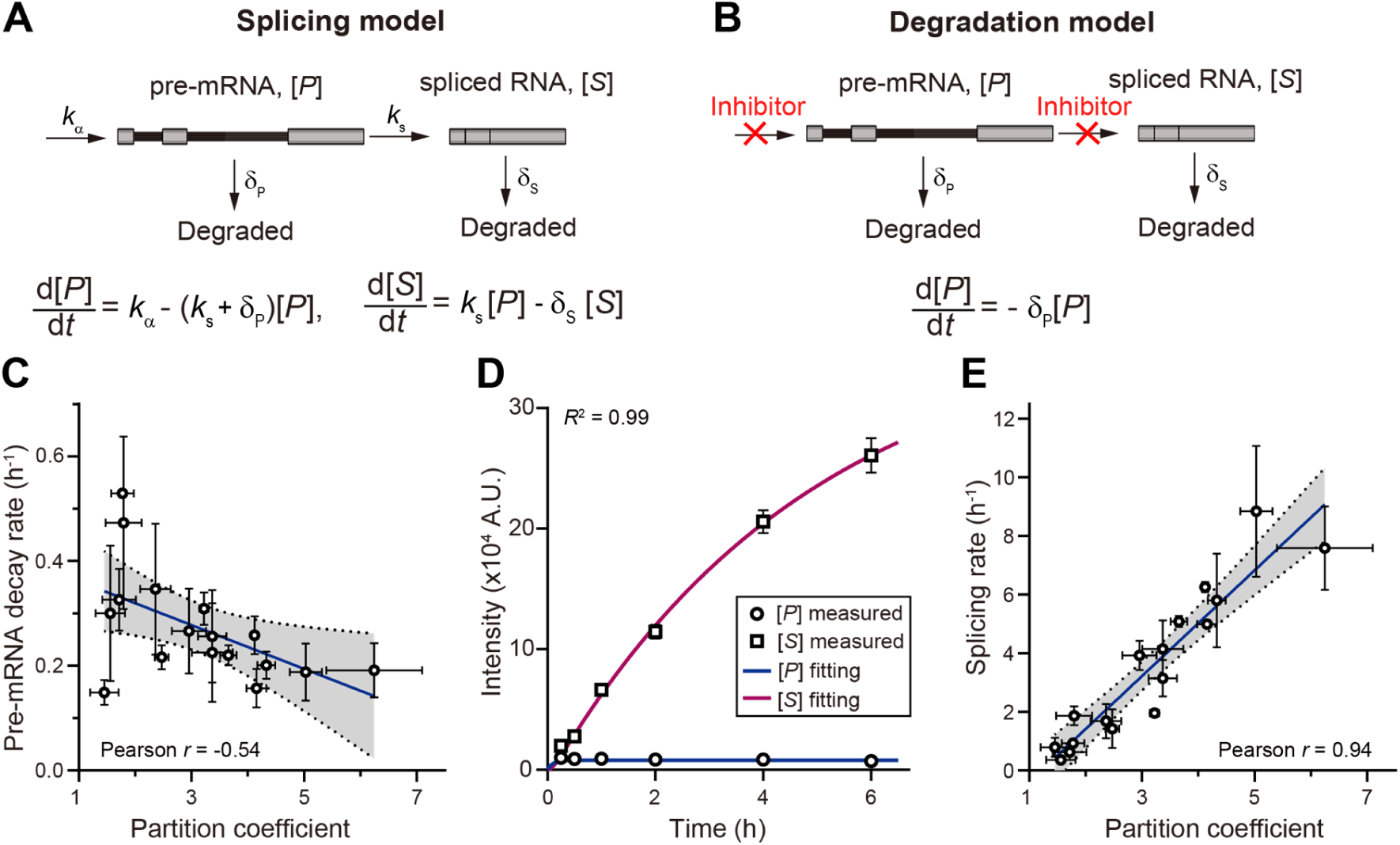
Nuclear speckle localization correlates with nuclear RNA degradation and splicing kinetics. (A–B) Schematic illustration of the assay for measuring splicing rate (A) and degradation rate (C) with the related mass-action equations. *k*_α_, *k*_s_, *δ*_p_, and *δ*_s_ represent the rate constant of transcription, splicing, pre-mRNA degradation, and spliced mRNA degradation, respectively. [*P*] and [*S*] represent the concentration of the pre-mRNA and spliced mRNA respectively. (D) Correlation between the pre-mRNA degradation rate and partition coefficient of pre-mRNA from minigenes. (E) Representative fitting of time-dependent change pre-mRNA concentration [*P*] and spliced mRNA concentration [*S*] for F_3’-SR-5’_. (F) Correlation between the splicing rate and partition coefficient of pre-mRNA from minigenes. In (C) and (E), error bars report the standard deviation from 2-7 biological replicates, with each replicate including 21–2008 cells. Pearson correlation coefficients are shown on the plot. In (D), error bars report the standard error of the mean calculated from single-cell measurements. Each data point includes 645-1095 cells. *R*^2^ from nonlinear regression is displayed on the plot.

We employed qPCR to measure the percentages of pre-mRNA and intron lariat that constitute the total intronic probe signal (Table S1). To ensure that the intronic and exonic fluorescent signals reflect the stoichiometry of the represented molecular abundance, we used a control sample treated with Plad B. Since transcripts are predominately unspliced upon Plad B treatment, the intron and exon abundances should be equal. We therefore adjusted the imaging parameter during data acquisition so that the fluorescent signals from the intronic and exonic probes were equal to stoichiometrically reflect the molecular abundance. The exact same imaging parameters were used in the kinetic measurements. To further constrain the parameters, we measured the pre-mRNA nuclear degradation rate of each construct by first inhibiting splicing with Plad B and then blocking transcription with triptolide.^38,39^ We then fixed and imaged cells at multiple time points and fitted the decay of pre-mRNA fluorescence intensity to an exponential function to determine degradation rates. Our result demonstrates a mild negative correlation between the speckle partition coefficient and pre-mRNA degradation rate, indicating that speckle localization protects RNA from nuclear degradation to some extent (Figure 7C, Table S4).

The kinetic model provided a good fit to the time-dependent changes measured in pre-mRNA and spliced mRNA levels (Figure 7D), allowing us to confidently assign a splicing rate to each of the constructs. Interestingly, we found that splicing rate shows close to linear relationship with the speckle partition coefficient (Figure 7E, Table S4). Importantly, this positive correlation is independent of splicing outcome. For instance, F_3’-SR-5’_ shows a high speckle localization propensity and a fast splicing rate with an exon included splicing outcome. hnRNP motif-containing minigenes demonstrate a moderate splicing rate, consistent with their moderate speckle localization propensity, with skipping of the entire hnRNP motif-containing region. It is insightful to compare these hnRNP motif-containing minigenes with F_SR_: even though the splicing outcome is identical in both cases (Figure 5C), F_SR_’s splicing rate is 8-fold slower than that of F_3’-HN-5’_, leading to mostly unspliced product, consistent with its much lower speckle localization. Remarkably, this difference in speckle localization was already apparent in the minimal constructs (Figure 1D and E), i.e., without any splicing context.

## Discussion

In this study, we took a synthetic approach to systematically identify the determinants of RNA speckle localization, providing new insights into speckle function. This approach avoided the confounding effects of genomic architecture and RNA processing. An interesting pattern emerges from our results: sequences that characterize unspliced pre-mRNA, namely, unspliced exon-like sequences and intron-like sequences, are also those promoting speckle localization. This pattern holds even for transcripts that do not undergo any splicing. We found that speckle localization is remarkably sensitive to sequence variation, with even single nucleotide disease-associated exonic variants affecting it. Finally, we showed that speckle localization is strongly correlated with splicing kinetics and splicing outcomes. Together, our results highlight nuclear speckles as an organelle attracting unspliced pre-mRNA and facilitating splicing.

The observed sequence determinants suggest that spliceosome assembly is tightly connected to speckle localization. To explore this connection further, we introduced sequence perturbations and knocked down spliceosomal proteins. We found that speckle localization of M_3’-SR-5’_ is impaired when 5’ splice site recognition by U1 snRNP is weakened. Splicing-related sequence elements around the 3’ splice site also contribute to speckle localization to various extents. Specifically, the polypyrimidine tract interaction with U2AF2 appears to contribute the most, followed by the 3’ splice site interaction with U2AF1. Also, treatment with the splicing inhibitor Pladienolide B had minimal effect on the partition coefficient of one tested construct (Figure S2E, Supplemental notes), consistent with our recent work showing that splicing inhibition using Plad B does not significantly affect speckle localization of pre-mRNA transcriptome-wide.^12^ Finally, it is interesting to note that M_3’-SR-5’_ has a much higher partition coefficient than M_3’-SR_ and M_SR-5’_, suggesting synergy between U1 snRNP and U2AF1/U2AF2 or other associated factors, and relating to the idea of exon definition.^40,41^ Together, these observations all point to the early spliceosome E complex as being associated with speckle localization.^42^

Our reporter-based results elucidate previous studies on endogenous genes. First, previous work has highlighted speckles as hubs of post-transcriptional splicing regulation.^43^ Indeed, one of the most striking features of speckle-localized transcripts is their high abundance of post-transcriptionally spliced introns: at least three in the majority of speckle-localized transcripts compared to zero or one in the majority of those not localizing to speckles.^12^ However, transcriptomic analysis struggled to establish causality due to confounding effect of genomic architecture (e.g., speckle localized transcripts also have high GC content and short introns) and the inability to fully decouple speckle localization from the splicing reaction.^12^ Our results indicate that the presence of features resembling unspliced pre-mRNA, such as unspliced exons and intronic sequences, serves as an indication of incomplete splicing and drives transcripts to speckles. Second, we observed that higher pre-mRNA speckle localization propensity couples with faster splicing rate, supporting the previous observation that nuclear speckles facilitate splicing of speckle-localized transcripts.^9,44^ Finally, we also observed that speckle localization provides a mild protective effect against nuclear degradation. This observation reflects a previous notion that a negative regulator of nuclear exosome is enriched in nuclear speckle.^45^ Collectively, our results suggest that beyond the impact of gene position, RNA localization to nuclear speckles is directly coupled to the splicing logic in RNA sequences and highlight the roles of nuclear speckles in splicing regulation of the human transcriptome.

Historically, RNA localization was considered in the context of asymmetric cytoplasmic distributions in highly polarized cells with specialized functions, such as oocytes or neurons. Our work contributes to an increasing body of literature that argues that RNA localization to membraneless organelles is a fundamental mechanism that regulates all aspects of RNA processing, from splicing to decay. Similar synthetic approaches can be applied to other membraneless organelles. For instance, transcriptomic studies have shown that P-bodies contain AU-rich transcripts^46^ whereas stress granule localization is associated with the presence of m^6^A modifications.^47^ Identifying their sequence specificities beyond such broad RNA content preferences could provide more insight into their function. We also envision our approach will find utility in the development of drugs for splicing-related diseases. As we demonstrated with branaplam, the effects of splice-modulating drugs can be quantified directly from FISH signal in a minimal system, without the need for fluorescent or other reporter assays.

### Limitations of the study

Even though our results generalize to disease-associated genes and are consistent with previous studies on endogenous genes, the use of reporter constructs limits our ability to capture the entire complexity of endogenous genes. In particular, we did not measure the localization behavior of many-exon genes. We are also unable to capture the effect of chromatin environment on speckle localization. Nevertheless, the principles identified here should guide further studies into the localization of endogenous genes.

The molecular mechanism behind the peripheral localization of hnRNP-containing transcripts (like M_HN_) is not clear; we speculate that it may be related to a reported depletion of hnRNP protein inside speckles^24,25^ and possibly to the intra-speckle organization of speckle-localized transcripts.^8^ It is also not clear what mechanism drives M_5’-HN_ transcripts so strongly to speckles compared to all other hnRNP motif containing constructs. One possible explanation is that this behavior is due to the nuclear cap-binding complex, which is known to facilitate association of U1 snRNP with nearby 5’ splice sites, as part of the definition of the first exon during early spliceosome assembly.^48^ Alternatively, a recent study revealed that an unpaired 5’ splice site followed by a downstream polyA sequence promotes nuclear RNA degradation.^49^ The high speckle enrichment of M_5’-HN_ transcripts may therefore also partially stem from a differential degradation rate in the nucleoplasm versus nuclear speckles, leading to a faster depletion of nucleoplasm-localized transcripts. These explanations, however, do not address the sensitivity of M_5’-HN_ partition coefficient to U2AF1 and U2AF2 knockdown.

## Acknowledgments

This project was supported by the NIH Director’s New Innovator Award (1DP2GM128185-01) to JF, a Simons Investigator Award and NSF MCB-2226731 to OR, NSF MCB-2246530 to JF and OR, a Yen Foundation Fellowship to XF and SP, and a Life Sciences Research Foundation Fellowship from Additional Ventures to SEL. We thank Jane Chen for help with the graphical abstract.

## Author contributions

Conceptualization: JF, OR, LW, MAA

Experiment and analysis: LW, MAA, XF, SP, SEL, MS

Supervision: JF, OR

Writing: JF, OR, LW, MAA

## Star methods

Key resources table.

| REAGENT or RESOURCE | SOURCE | IDENTIFIER |
|---|---|---|
| Antibodies |  |  |
| Secondary antibody against mouse | Jackson ImmunoResearch | #715-005-150 |
| Secondary antibody against rabbit | Jackson ImmunoResearch | #711-005-152 |
| Mouse antibody against SRRM2 | Sigma Aldrich | #S4045 |
| Mouse antibody against SRSF1 | Invitrogen | #32-4600 |
| Rabbit antibody against SON | Novus | #NBP2-55411 |
| Rabbit antibody against hnRNPA1 | Abcam | #ab177152 |
| Mouse antibody against U1A | Santa Cruz | #sc-376027 |
| Rabbit antibody against U2AF1 | Abcam | #ab172614 |
| Rabbit antibody against U2AF2 | Abcam | #ab177152 |
| Chemicals, peptides, and recombinant proteins |  |  |
| BsaI | New England Biolabs | #R3733 |
| BsmBI | New England Biolabs | #R0739 |
| MfeI-HF | New England Biolabs | #R3589S |
| PciI | New England Biolabs | #R0665S |
| Dulbecco's Modified Eagle Medium | Gibco | #11965092 |
| R&D Fetal Bovine Serum - TET Tested | R&D Systems | #S10350 |
| Tet system approved fetal bovine serum | Gibco | #631105 |
| Sodium pyruvate | Thermo Fisher | #11360070 |
| Penicillin-streptomycin | Thermo Fisher | #15070063 |
| Lipofectamine 2000 reagent kit | Invitrogen | #11668027 |
| Lipofectamine 3000 reagent kit | Invitrogen | #L3000001 |
| Opti-MEM | Thermo Fisher | #31985070 |
| Doxycycline | Santa Cruz | #sc-204734B |
| Pladienolide B | Cayman Chemicals | #445493-23-2 |
| Triptolide | Sigma Aldrich | #T3652-1MG |
| Branaplam | MedChemExpress | #HY-19620 |
| TURBO DNase | Thermo Fisher | #AM2238 |
| Trizol | Thermo Fisher | #15596026 |
| Phenol/chloroform | Thermo Fisher | #AM9720 |
| Lipofectamine RNAiMax reagent | Invitrogen | #13778150 |
| Amino-11-ddUTP | Lumiprobe | #15040 |
| Terminal transferase | New England Biolabs | #M0315L |
| Alexa Fluor 647 succinimidyl ester | Invitrogen | #A20106 |
| CF568 succinimidyl ester | Sigma Aldrich | #SCJ4600027 |
| Alexa Fluor 488 succinimidyl ester | Invitrogen | #A20000 |
| Paraformaldehyde | Electron Microscopy Sciences | #15710 |
| Triton X-100 | Thermo Fisher | #BP151-100 |
| Vanadyl ribonucleoside complexes | Sigma-Aldrich | #R3380 |
| Formamide | Ambion | #AM9342 |
| Dextran sulphate | Sigma-Aldrich | #D8906 |
| Dithiothreitol | Sigma-Aldrich | #10197777001 |
| Ultrapure BSA | Invitrogen | #AM2618 |
| DAPI | Thermo Fisher | #62248 |
| Glucose oxidase | Sigma-Aldrich | #G2133 |
| Catalase | Sigma-Aldrich | #C3515 |
| Critical commercial assays |  |  |
| RNA extraction kit | QIAgen | #74104 |
| RNA extraction kit | QIAgen | #75144 |
| Phusion High Fidelity kit | New England Biolabs | #E0553L |
| Reverse Transcription SuperScript IV | Thermo Fisher | #18090010 |
| GeneJET PCR Purification Kit | Thermo Fisher | #K0701 |
| Neon<sup>TM</sup> Transfection System | Thermo Fisher | #MPK5000 |
| iScript cDNA synthesis kit | Bio-Rad | #1708890 |
| SsoAdvanced Universal SYBR Green Supermix | Bio-Rad | #1725272 |
| LightCycler 480 SYBR Green I Master mix | Roche | #04887352001 |
| Deposited data |  |  |
| Code used for quantification of partition coefficients, splicing rates, and decay rates | This paper | (https://github.com/JingyiFeiLab/2025_NS_localization ) |
| Plasmids | This paper | Addgene |
| Experimental models: Cell lines |  |  |
| HeLa Tet-On cells | TaKaRa Bio, USA | #631183 |
| Recombinant DNA |  |  |
| Plasmid: pU1_000 | Hatch et al.<sup>50</sup> | Addgene #186773 |
| Plasmids: Minimal: M<sub>SR</sub> M<sub>5'-SR</sub> M<sub>SR-3'</sub> M<sub>3'-SR</sub> M<sub>SR-5'</sub> M<sub>3'-SR-5'</sub> M<sub>HN</sub> M<sub>5'-HN</sub> M<sub>HN-3'</sub> M<sub>3'-HN</sub> M<sub>HN-5'</sub> M<sub>3'-HN-5'</sub> M<sub>5'-HN,upstream cryptic AG removed</sub> M<sub>5'-HN,upstream cryptic PY removed</sub> M<sub>3'-SR-weak5'</sub> M<sub>noBP,3'-SR-5'</sub> M<sub>weakPY,3'-SR-5'</sub> M<sub>noAG,3'-SR-5'</sub> M<sub>A1</sub> M<sub>A2</sub> M<sub>A3</sub> M<sub>A4</sub> M<sub>A5</sub> M<sub>A6</sub> M<sub>B1</sub> M<sub>B2</sub> M<sub>B3</sub> M<sub>B4</sub> M<sub>B5</sub> M<sub>B6</sub> M<sub>3'-SR-5'-SR</sub> M<sub>3'-HN-5'-HN</sub> M<sub>3'-SR-5'-HN</sub> M<sub>3'-SR-HN</sub> M<sub>SR-5'-HN</sub> M<sub>SR-HN</sub> M<sub>3'-HN-5'-SR</sub> M<sub>HN-5'-SR</sub> M<sub>3'-HN-SR</sub> M<sub>HN-SR</sub> M<sub>SMN1</sub> M<sub>SMN2</sub> M<sub>COLQ-WT</sub> M<sub>COLQ-MUT</sub> Minigene: F<sub>3'-SR-5'</sub> F<sub>SR-5'</sub> F<sub>3'-SR</sub> F<sub>SR</sub> F<sub>3'-HN-5'</sub> F<sub>HN-5'</sub> F<sub>3'-HN</sub> F<sub>HN</sub> F<sub>3'-SR-5'-HN</sub> F<sub>3'-SR-HN</sub> F<sub>SR-5'-HN</sub> F<sub>SR-HN</sub> F<sub>3'-HN-5'-SR</sub> F<sub>HN-5'-SR</sub> F<sub>3'-HN-SR</sub> F<sub>HN-SR</sub> F<sub>3'-N-5'-HN</sub> F<sub>N-5'-HN</sub> F<sub>3'-N-5'-SR</sub> F<sub>N-5'-SR</sub> F<sub>A1</sub> F<sub>A2</sub> F<sub>A3</sub> F<sub>A4</sub> F<sub>A5</sub> F<sub>A6</sub> F<sub>B1</sub> F<sub>B2</sub> F<sub>B3</sub> F<sub>B4</sub> F<sub>B5</sub> F<sub>B6</sub> | This study | N/A |
| Plasmids: U1<sub>weak</sub> U1<sub>WT</sub> | This study | N/A |
| Software and algorithms |  |  |
| Stellaris Probe Designer | LGC Biosearch Technologies | https://www.biosearchtech.com/support/tools/design-software/stellaris-probe-designer |
| Fiji (ImageJ) | Schindelin, J., Arganda-Carreras, I., Frise, E. et al.<sup>51</sup> | https://imagej.nih.gov/ij/ |
| NIS-Elements | Nikon | https://www.microscope.healthcare.nikon.com/products/software/nis-elements |
| GraphPad Prism 8 | GraphPad Software | https://www.graphpad.com |
| MATLAB | MathWorks | https://www.mathworks.com/products/matlab.html |
| Cellpose | Stringer, C., Wang, T., Michaelos, M. et al.<sup>52,53</sup> | https://github.com/MouseLand/cellpose |
| BLAST | National Library of Medicine | https://blast.ncbi.nlm.nih.gov/Blast.cgi |
| Snapgene v7.0.1 | Dotmatics | https://www.snapgene.com/ |
| Other |  |  |
| 8-well imaging chamber, #1.5 cover glass | Cellvis | #C8-1.5H-N |
| 96-well imaging chamber, #1.5 cover glass | Cellvis | #P96-1.5H-N |
| P-6 Micro Bio-Spin Column | Bio-Rad | #7326222 |

## Method details

### Plasmid design and construction

Plasmid design was based on our earlier work.^8^ Briefly, 248bp SR-enriched and hnRNP-enriched sequences were designed by randomly mutating initial sequences containing repeats of validated binding motifs. Sequences were verified to not contain any predicted splice site sequence and to keep a similar enrichment of SR or hnRNP motifs. To avoid a long repetition, the second enriched regions in the dual constructs were designed separately (and are also longer at 392bp). Fragments were then synthesized (Integrated DNA Technologies) and assembled using a Golden Gate assembly protocol into a backbone containing a Tet-responsive promoter and the SV40 polyA sequence in the strong orientation. Minigene constructs (the F series) were generated using a backbone containing, in addition, the Chinese hamster *DHFR* exon 1 and intron 1, an intronic sequence derived from *DHFR* intron 3, and finally, the concatenation of *DHFR* exons 4 through 6. U1 plasmids were modified from pU1_000, which was a gift from Eugene Yeo (Addgene plasmid #186773). All plasmids were verified using whole-plasmid sequencing (Plasmidsaurus). Sequence information can be found in Supplementary Data S1.

### Cell culture

HeLa Tet-On cells (TaKaRa) were cultured in Dulbecco’s Modified Eagle Medium (DMEM, Gibco) containing 4.5 g/L glucose. The medium was supplemented with 10% fetal bovine serum (FBS, Gibco), 1 mM sodium pyruvate (Gibco), and 50 U/mL penicillin-streptomycin (Gibco). Regular testing for mycoplasma contamination was performed. For microscopy-based assays, cells were seeded into imaging-compatible 8-well (Cellvis, C8-1.5H-N) or 96-well chamber (Cellvis, P96-1.5H-N). For quantification of intron-exon junction to intron ratio by qPCR, 24-well plates (Corning, 353047) were used. For other biochemical assays, 12 well dishes (Corning, #3513) were used. Experiments were performed with cells at 70–80% confluency.

### Transfection and drug treatments

Circular plasmids were transfected using Lipofectamine 3000 reagent (Invitrogen) following the manufacturer’s protocol. For each well of an 8-well chamber, the transfection mixture was prepared with 30 μL reduced serum medium (opti-MEM, Thermofisher), 0.4 μL P3000 reagent, 200 ng plasmid DNA, and 0.6 μL Lipofectamine 3000 reagent. Before transfection, Tet-free DMEM medium (DMEM with 10% Tet system approved FBS (TaKaRa, 631105)) was added to replace the culture medium. The DNA–lipid complexes were subsequently dispensed into each well. The transfection medium was replaced with fresh Tet-free DMEM at 6 h post transfection. For chambers of different sizes, reagent volumes were adjusted proportionally based on the surface area.

For linearization, circular plasmids were digested by MfeI-HF(NEB, R3589S) or PciI (NEB, R0665S) and then purified by the GeneJET PCR Purification Kit (Thermofisher, K0701). Digestion efficiency was verified by agarose gel electrophoresis. Due to the reduced transfection efficiency of linearized plasmids using Lipofectamine, electroporation was employed instead. Electroporation was performed using the Neon™ Transfection System (Thermofisher, MPK5000) according to the manufacturer’s protocol. For each well of an 8-well chamber, 1.5 × 10⁵ cells were electroporated with 500 ng of linearized plasmid DNA using a 10 μL Neon tip. Electroporated cells were plated in Tet-free DMEM and incubated overnight.

Induction, splicing inhibition, and branaplam treatment were performed 24 h after transfection in Tet-free DMEM using the following conditions: doxycycline (2 μg/mL, Santa Cruz Biotechnology, sc-204734B) for 2 h; Pladienolide B (Plad B, 100 nM, Cayman Chemical, 445493-23-2) for 4 h; and branaplam (MedChemExpress, HY-19620) at final concentrations of 20 nM, 50 nM, or 100 nM, each applied for 4 h.

### siRNA-mediated knockdown

U1A, U2AF1, U2AF2, and SRSF1 knockdown was performed using Lipofectamine RNAiMax reagent (Invitrogen, 13778030) following the manufacturer’s protocol. DsiRNAs targeting U1A (hs.Ri.SNRPA.13.2), U2AF1 (hs.Ri.U2AF1.13.2), U2AF2 (hs.Ri.U2AF2.13.2), SRSF1 (hs.Ri.SRSF1.13.2), as well as a scrambled negative control DsiRNA (51-01-19-08) were purchased from IDT. For each well of an 8-well chamber, the siRNA-lipid complex was prepared by combining 25 μL Opti-MEM, 0.75 μL Lipofectamine RNAiMax reagent, and 0.25 μL siRNA (10 μM). Plasmid transfection was performed 24 h after siRNA-mediated knockdown.

Knockdown of hnRNPA1 was carried out via electroporation using the Neon™ Transfection System (Thermo Fisher, MPK5000), following the manufacturer’s instructions. For each well of an 8-well chamber, 1.5 × 10⁵ cells were electroporated with 1.6 μL of a combined siRNA mixture targeting hnRNPA1(hs.Ri.HNRNPA1.13.1 and hs.Ri.HNRNPA1.13.2, IDT, 30 μM each) using a 10 μL Neon tip. For negative control, cells were electroporated with an equivalent concentration of scrambled control siRNA. After electroporation, cells were seeded in Tet-free DMEM and allowed to recover for 24 h before plasmid transfection.

### Long-read Nanopore sequencing

Nanopore sequencing was performed with MinION Mk1B devices using the PCR-cDNA Barcoding Kit (SQK-PCB114.24) and the R10.4.1 flow cells according to the manufacturer’s protocol. The protocol selects for polyadenylated transcripts by ligating an adapter at the 3’OH end of the RNA molecule and performs strand switching at the 5’ end. Samples were barcoded using the provided adapters through 14 to 16 PCR cycles and pooled together in batches of 16. Subsequent sequencing provided complete transcript sequence. MinKNOW software version 24.11.10 was used to produce pod5 files.

These files were then basecalled using dorado version 0.8.2 using the super accuracy model dna_r10.4.1_e8.2_400bps_sup@v5.0.0 and the ––no-trim option on one V100 GPU. The resulting BAM file was demultiplexed using dorado with options ––kit-name SQK-PCB114-24 ––no-trim. After converting the demultiplexed BAM files to FASTQ format, pychopper was used to orient the reads using option –k PCS111. Finally, oriented reads were aligned to the plasmid sequences using minimap2 with options –x splice ––cs –uf –k14. Aligned reads were then visualized, manually inspected, and summarized using custom scripts. Reads aligning in the reverse strand were rare in all constructs and were ignored in the analysis since they are not predicted to bind to any of our probes.

### RT-PCR for biochemical assays

Cells were induced 22 h after transfection, followed 2 h later by RNA extraction (QIAgen RNeasy minikit, 74116) using a QIAcube following the manufacturer’s protocol. DNA was removed using TURBO DNase (ThermoFisher, AM1907) in a 30 μL reaction. RNA concentration was then adjusted to 100 ng/μL using NanoDrop One. For reverse transcription, 200 ng of RNA was used for a 10 μL reaction using SuperScript IV (ThermoFisher, 18090010) with gene-specific primers at 100 nM concentration for 1 h, according to the manufacturer’s protocol. PCR reactions were carried out in an Eppendorf X50S 96-well thermocycler using Phusion High Fidelity polymerase (New England Biolabs, M0530L). The reverse transcription product was diluted 5-fold with water and 2.4 μL was used in a 30 μL PCR reaction according to the manufacturer’s instructions. PCR was run for 21 or 22 cycles allowing 2 min for extension. For each sample, 25 μL was loaded in a 0.7 cm 1.5% agarose gel and quantified in an iBright gel imager after 30 min of post-staining with ethidium bromide and three 10 min washes for destaining on an orbital shaker.

### Quantitative real-time PCR

#### Quantification of exon-intron junction to intron ratio

RNA was extracted after 2 h transcription induction using Trizol reagent (ThermoFisher, 15596026), followed by a TURBO DNase (ThermoFisher, AM2238) treatment and a phenol/chloroform (ThermoFisher, AM9720) extraction. cDNA was synthesized from 100 ng of total RNA in a reaction mixture containing 250 nM of a construct-specific reverse primer (Table S5), 10% DMSO (ThermoFisher, BP231), 1 mM dNTPs (NEB, N0447S), and 10 mM DTT (Sigma–Aldrich, 10197777001). Reverse transcriptase from iScript cDNA synthesis kit (Bio-Rad, 1708890) was diluted 1:20 and added to the mixture. The RT reaction was performed according to the manufacturer’s instructions. qPCR was performed on a CFX real-time PCR system (Bio-Rad) using primer pairs listed in Table S5. Each 20 μL reaction contained 10 μL of 2× SYBR Green Supermix (Bio-Rad, 1725272), 1 μL of cDNA, 2 μL of forward and reverse primers (2.5 μM each), and 7 μL of nuclease-free water. The following cycling parameters were used: 95 °C for 30 s (initial denaturation), followed by 40 amplification cycles at 95 °C for 10 s and 60 °C for 30 s. Ct values obtained from primer sets targeting the intron and the exon-intron junction were used to calculate relative ratios. To account for potential differences in primer efficiency, Ct values were also measured in a control sample treated with Plad B, in which transcripts are predominantly in the pre-mRNA form. The difference in efficiency between primer sets was quantified based on this control and used to correct the calculated ratios.

#### Quantification of PSI

Reverse transcription was performed as in the RT-PCR for biochemical assays described above. In addition, RNA from cells transfected using constructs producing a unique isoform (either the exon inclusion or the exon skipping isoforms) were serially diluted with RNA from untransfected cells and processed with the other samples to assess qPCR efficiency.

qPCR was performed on a LightCycler 480 II system (Roche) using primer pairs listed in Table S5. Each 10 μL reaction contained 5 μL of 2X LightCycler 480 SYBR Green I Master mix (Roche, 04887352001), 1 μl of diluted cDNA, primers at a final concentration of 400 nM and nuclease-free water. The following cycling parameters were used: 95 °C for 30 s (initial denaturation), followed by 40 amplification cycles at 95 °C for 10 s and 60 °C for 30 s. The measured qPCR efficiency and Ct values were used to calculate the relative amounts of each species and the corresponding PSI for each sample.

### Assay for measurement of splicing and degradation rates

For measuring pre-mRNA degradation rates, RNA splicing was inhibited by treating cells with Plad B for 2 h at 24 h post-transfection, followed by a 2 h induction with doxycycline in the continued presence of Plad B. Subsequently, transcription was blocked by adding 4 μM Triptolide (Sigma-Aldrich, T3652) for different durations depending on the degradation kinetics of the constructs: 0, 1, 2, and 3 h for slowly degraded constructs, and 0, 20, and 40 min, and 1 h for rapidly degraded constructs. After treatment, cells were fixed, and RNA FISH was performed using intron targeting probes. For measuring splicing rates, doxycycline was added at 24 h post-transfection to induce transcription for 15 min, 30 min, 1 h, 2 h, 4 h, and 6 h. Following treatment, cells were fixed, and RNA FISH was performed using both intron and exon targeting probes. Fluorescence images across time-course samples were recorded using automated multi-well imaging on a 96-well platform.

### Fluorescence labeling of FISH probes and secondary antibodies

FISH probes were designed using the Stellaris Probe Designer (Biosearch Technologies) and purchased from IDT. Amino–dideoxyuridine triphosphate (ddUTP) was conjugated to the 3′ end of each oligonucleotide using a terminal transferase reaction, as previously described.^54^ In a 30 μL reaction, 20 μL of pooled oligonucleotides (100 μM) were combined with 6 μL of 1 mM Amino-11-ddUTP (Lumiprobe, 15040), 1.2 μL of terminal deoxynucleotidyl transferase (NEB, M0315L) and 3 μL of 10x terminal transferase reaction buffer (NEB, M0315L) and incubated overnight at 37 °C. For fluorophore conjugation, the modified probes were first purified using a P-6 Micro Bio-Spin Column (Bio-Rad, 7326222), and then incubated overnight at 37 °C with 25 μg of Alexa Fluor 647 NHS Ester (Invitrogen, A20106) or CF568 succinimidyl ester (Sigma Aldrich, SCJ4600027) in 0.1 M sodium bicarbonate (pH 8.5). Labeled probes were purified first by ethanol precipitation and then with a P-6 Micro Bio-Spin column. All probes were labeled with an efficiency above 75%, as quantified by UV-Vis spectrums. The exact sequences of the FISH probes are provided in Table S5.

To label secondary antibodies, 24 μL of Donkey Anti-Mouse IgG (Jackson ImmunoResearch, 715-005-150) or Donkey Anti-Rabbit IgG (Jackson ImmunoResearch, 711-005-152) at 1 mg/mL was mixed with 3 μL of 1 M sodium bicarbonate (pH 8.5) and 3 μL of 10× PBS. Alexa Fluor 488 NHS Ester (Invitrogen, A20000) or CF568 succinimidyl ester (Sigma Aldrich, SCJ4600027) were added in amounts ranging from 1 to 3 μg. The reaction was incubated at room temperature for 1 h to allow conjugation. After labeling, unreacted dye was removed using a P-6 Micro Bio-Spin Column.

### RNA fluorescence in situ hybridization

RNA FISH and immunostaining were performed according to a previously published protocol.^55^ Cells were fixed at room temperature for 10 min using 4% paraformaldehyde (PFA; Electron Microscopy Sciences, 15710) in 1× PBS. Following fixation, three washes with 1× PBS were performed. Cells were permeabilized on ice for 10 min with 0.5% Triton X-100 (Thermo Fisher, BP151-100) and 2 mM vanadyl ribonucleoside complexes (Sigma-Aldrich, R3380) in 1× PBS. Cells were subsequently rinsed twice with 1× PBS and twice with 2x saline-sodium citrate (SSC), followed by a final wash in FISH wash solution containing 10% formamide (Ambion, AM9342) in 2× SSC. Hybridization was performed overnight at 37 °C in the dark by adding probe-containing hybridization buffer [FISH wash solution supplemented with 5 nM of each fluorescently labeled probe, 10% dextran sulfate (Sigma-Aldrich, D8906-10g) and 10 mM dithiothreitol (Sigma-Aldrich, 10197777001)]. The following day, excess probes were removed by washing the cells with FISH wash solution for 30 minutes at 37 °C.

### Immunofluorescence staining

To prevent probe dissociation during immunostaining, post-fixation was performed. Afterward, cells were washed with 1x PBS and incubated in a blocking buffer containing 0.1% ultrapure BSA (Invitrogen, AM2618) in 1x PBS for 30 min at room temperature. Primary antibodies were diluted in the same blocking solution using the following ratios: mouse anti-SRRM2 (1:2000, Sigma Aldrich, S4045), rabbit anti-SON (1:200, Novus, NBP2-55411), mouse anti-U1A (1:100, Santa Cruz, sc-376027), rabbit anti-U2AF1 (1:100, Abcam, ab172614), rabbit anti-U2AF2 (1:1000, Abcam, ab37530), mouse anti-SRSF1 (1:250, Invitrogen, 32-4600) and rabbit anti-hnRNPA1 (1:250, Abcam, ab177152). Antibody solutions were added to each well and allowed to incubate at room temperature for 1 h, followed by three washes with 1x PBS. Subsequently, fluorescently labeled secondary antibodies (1:200 dilution in blocking solution) were applied and incubated for 1 h at room temperature. Afterward, cells were washed three more times with 1x PBS, and then stained with DAPI (Thermofisher, 62248) in 1x PBS. Cells were stored in 4x SSC at 4 °C until imaging.

### Image acquisition

Fluorescence imaging was performed using a Nikon Ti2-E inverted confocal microscope (Nikon AX-R) with GaAsP PMT detectors (DUX-ST, Nikon) operating in resonant scanning mode. A Plan Fluor 20× air objective (NA 0.50) was used for kinetic measurements, while a CFI Plan Apo 60× oil immersion objective (NA 1.40) was employed for all other imaging experiments. Excitation was performed by a multi-line laser unit (AS405/488/561/640, LUA-S4, Nikon) with appropriate laser and filter configurations. A step size of 0.3 μm was used for Z-stack acquisition. The pinhole was set to 2 Airy units throughout image acquisition. For high throughput acquisition, multi-well automated imaging was implemented using the JOBS module in Nikon NIS-Elements C software. An imaging buffer containing 50 mM Tris-HCl (pH 8), 10% glucose, 0.5 mg/mL glucose oxidase (Sigma-Aldrich, G2133), and 67 μg/mL catalase (Sigma-Aldrich, C3515) in 2x SSC was used to reduce photobleaching.

## Image analysis

### Quantification of partition coefficients

Individual cells were segmented with a custom-trained Cellpose model.^52,53^ The following image analysis was performed in MATLAB R2022b. Raw microscopy images were first imported using the Bio-Formats Importer.^56^ Based on the cell masks generated from Cellpose, individual region of interest (ROI) containing one segmented cell was cropped for downstream analysis. To determine the best focal plane for each cell, the Z-slice with the highest absolute Laplacian value was selected. Nucleus segmentation was carried out using Otsu’s algorithm on the DAPI channel. Transfected cells were subsequently determined based on global thresholding of RNA fluorescence intensity. To segment nuclear speckles, the intensities of SRRM2 or SON channels were first normalized across cells. A difference of Gaussians (DoG) filter was then used to subtract background, followed by global thresholding to detect speckles. This threshold was determined by the statistical properties (mean and standard deviation) of the DoG-filtered image. To ensure segmentation quality, cells with fewer than 5 detected nuclear speckles, or speckles smaller than 3 pixels in size, were excluded from further analysis. Finally, single-cell partition coefficients were calculated by dividing the mean RNA fluorescence intensity within nuclear speckles by the corresponding mean intensity in the surrounding nucleoplasm.

### Quantification of splicing and decay rates

A separate custom Cellpose model was trained and used to account for changes in imaging conditions due to the use of a different objective lens. The image analysis workflow followed the same procedure as described above, except nuclear speckle segmentation was omitted. For pre-mRNA decay rate quantification, the natural logarithm of fluorescence intensities obtained from intronic probes integrated over the entire cell was plotted as a function of time, and the degradation rate was estimated from the slope of the resulting linear regression.

For splicing rate quantification, single-cell fluorescence intensities integrated over the entire cell from intronic and exonic probes were extracted and averaged across cells to compute population-level means at each time point. Based on these measurements, the relative abundances of pre-mRNA [*P*] and spliced RNA [*S*] were estimated in units of fluorescence intensity using the following equations:

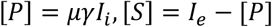

Here, *I*_i_ and *I*_e_ represent the single-cell mean intensities from intronic and exonic probes, respectively. The parameter μ denotes the pre-mRNA fraction among intronic signals, determined by qPCR (described above), while γ is a calibration factor that accounts for differences in intronic and exonic channels obtained by a sample treated with Plad B. Since imaging parameters were adjusted such that, in the sample treated by Plad B, intronic and exonic signals were approximately equal in intensity, the fitted value of γ was found to be close to 1.

The changes in their relative abundances over time were fitted using the following analytical expressions:

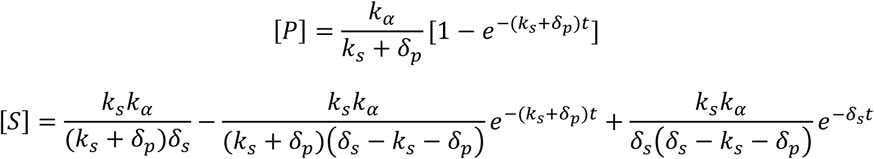

where *k*_a_ denotes the transcription initiation rate, *k*_s_ denotes the splicing rate, and δ_p_ and δ_s_denote degradation rates of pre-mRNA and spliced RNA, respectively. Here, the pre-mRNA degradation rate δ_p_ was fixed based on experimentally measured values as described above, while the remaining parameters were treated as free variables and estimated through fitting.

## Resource availability

### Lead contact

Further information and requests for resources and reagents should be directed to and will be fulfilled by the Lead Contact, JF

### Materials availability

All plasmids used in this study are available through the lead contact and will be made available through AddGene.

### Data and code availability

- The authors declare that all data supporting the findings of the present study are available in the article and its supplemental figures and tables, or from the corresponding author upon request.
- Code used for nuclear speckle partition coefficient analysis is available on GitHub (https://github.com/JingyiFeiLab/2025_NS_localization).

## Supplemental information (7)

Supplemental File: Supplemental notes, Figures S1-S7.

Supplemental Table S1

Supplemental Table S2

Supplemental Table S3

Supplemental Table S4

Supplemental Table S5

Supplementary Data S1

