## Supplemental information for "RNA localization to nuclear speckles follows splicing logic"

### Supplemental notes

#### Validations of constructs containing unintended transcripts

To ensure that RNA transcribed from our constructs does not deviate from the intended sequence, we performed extensive Nanopore full-length RNA sequencing. Importantly, we applied transcriptome-wide sequencing without using any gene-specific primers, ensuring that even transcripts significantly deviating from the intended sequence will be captured. We found that in almost all our constructs, the vast majority of sequenced transcripts are as intended (Figure S1). In some cases, however, we did identify a small fraction of unintended transcripts. These constructs are typically spliced, and as a result, we expect them to be efficiently exported from the nucleus, and therefore not affect our FISH measurements. Nevertheless, to ensure that these transcripts do not affect our imaging results, we validated them in a case-specific manner, as described below.

*Cryptic splicing in $M_{HN\text{-}3'}$*

The only construct showing high levels of unintended products is $M_{HN\text{-}3'}$. There, ~90% of reads were cryptically spliced due to the presence of weak cryptic 5' splice sites as determined by Nanopore sequencing (Figure S1B and E). Since short transcripts might be over-represented in Nanopore cDNA sequencing,[1,2] we further used gel electrophoresis analysis to quantify the fraction of spliced products, finding it to be ~50% (Figure S2C). Since only 1 out of 7 FISH probes is predicted to bind to the cryptic spliced product (Figure S2B, II), we expect our FISH imaging to predominately report on intended, unspliced transcripts, and hence not be affected by the spliced transcripts. To further test this, we performed FISH imaging on $M_{HN\text{-}3'}$ in the presence of the splicing inhibitor Plad B. Nanopore sequencing identified only <10% spliced reads, confirming efficient splicing inhibition (Figure S1E). We also observed an increase in the RNA FISH signal (Figure S2D), consistent with an increase in the unspliced transcript fraction, but no change in the partition coefficient (Figure S2E), supporting that the reported partition coefficient is specific to unspliced $M_{HN\text{-}3'}$ transcripts.

*Transcription read-through in $M_{5'\text{-}HN}$*

Unpaired 5' splice sites have been reported to suppress polyadenylation.[3] We indeed observed low levels of transcription read-through beyond the SV40 polyadenylation site in constructs carrying unpaired 5' spice sites. These read-through transcripts further underwent splicing using cryptic 3’ splice sites located downstream in the plasmid backbone. $M_{5'\text{-}HN}$ showed one of the highest percentages of read-through transcripts (~10%) (Figure S1C and E). To rule out the possibility that its high speckle localization is due to transcription read-through and subsequent cryptic splicing, we linearized the plasmid using one of two restriction enzymes targeting the polyadenylation site (Figure S2B, IV). We then transfected these linearized plasmids, and quantified the partition coefficient of the resulting transcripts, which are by design devoid of any read-through transcripts. Our data showed that these transcripts have a similar partition coefficient to the original circular plasmid (Figure S2F), confirming that the small contamination by read-through transcripts does not affect our results.

*Cryptic transcription in SMN1*

Most commonly used plasmid backbones are known to contain a cryptic eukaryotic transcription site inside the bacterial origin of replication sequence.[4] Indeed, in most of our constructs, we found a small (typically ~5%) fraction of transcripts originating from that site. These transcripts splice into our region of interest, and can therefore potentially bind a subset of our FISH probes. This is the case for the SMN1 and SMN2 constructs, with the former reproducibly exhibiting ~10% cryptic transcripts whereas the latter exhibiting only ~5% (Figure S1D and E). To exclude the possibility that the observed mildly higher partition coefficient of SMN1 compared to SMN2 was due to the additional cryptic transcripts in the former, we labeled two sets of FISH probes, one with AF647 (targeting an upstream region not present in the cryptic transcripts) and the other with CF568 (targeting a downstream region present in both intended and cryptic transcripts) (Figure S2B, III). We then performed two-color FISH imaging on uninduced cells, in which cryptic transcripts are expected to still be generated, since they are not driven by a dox-inducible promoter (Figure S2G). No FISH signal was detected from the AF647-labeled probes in the absence of doxycycline, suggesting that cryptic transcription has a negligible effect on our measurements.

### Supplemental Figures

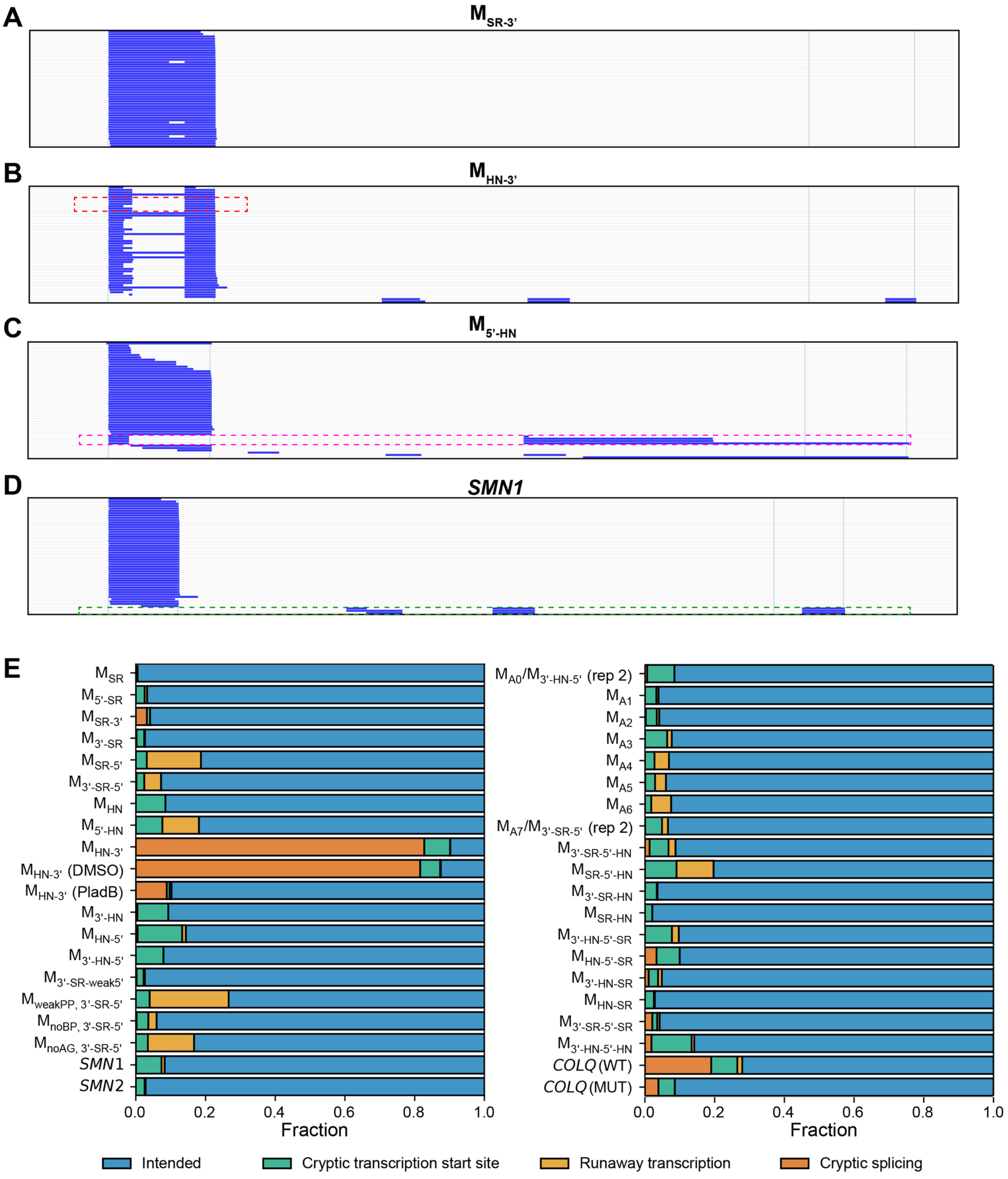


**Figure S1. Summary statistics of minimal constructs Nanopore sequencing.**
(A-D) Alignments of a random subset of long Nanopore reads from several minimal constructs. To allow simultaneous visualization of reads throughout the circular plasmid, the x-axis covers part of the plasmid twice. In particular, the intended transcript region (bounded by two thin

vertical lines) appears twice in each panel, once on the left and once on the right. While reads from some constructs are mostly as intended (A), others show cryptic splicing (B), transcription read-through (C), or cryptic transcript start (D), as highlighted by the dashed boxes.

(E) Summary statistics of read types in each sample. 5'-truncated and 3'-truncated reads were ignored. Such reads were generally a small minority of all reads. Moreover, they were also present in endogenous transcripts (*GAPDH*) and therefore likely represent technical artifacts due to RNA degradation during sample preparation or due to Nanopore sequencing.

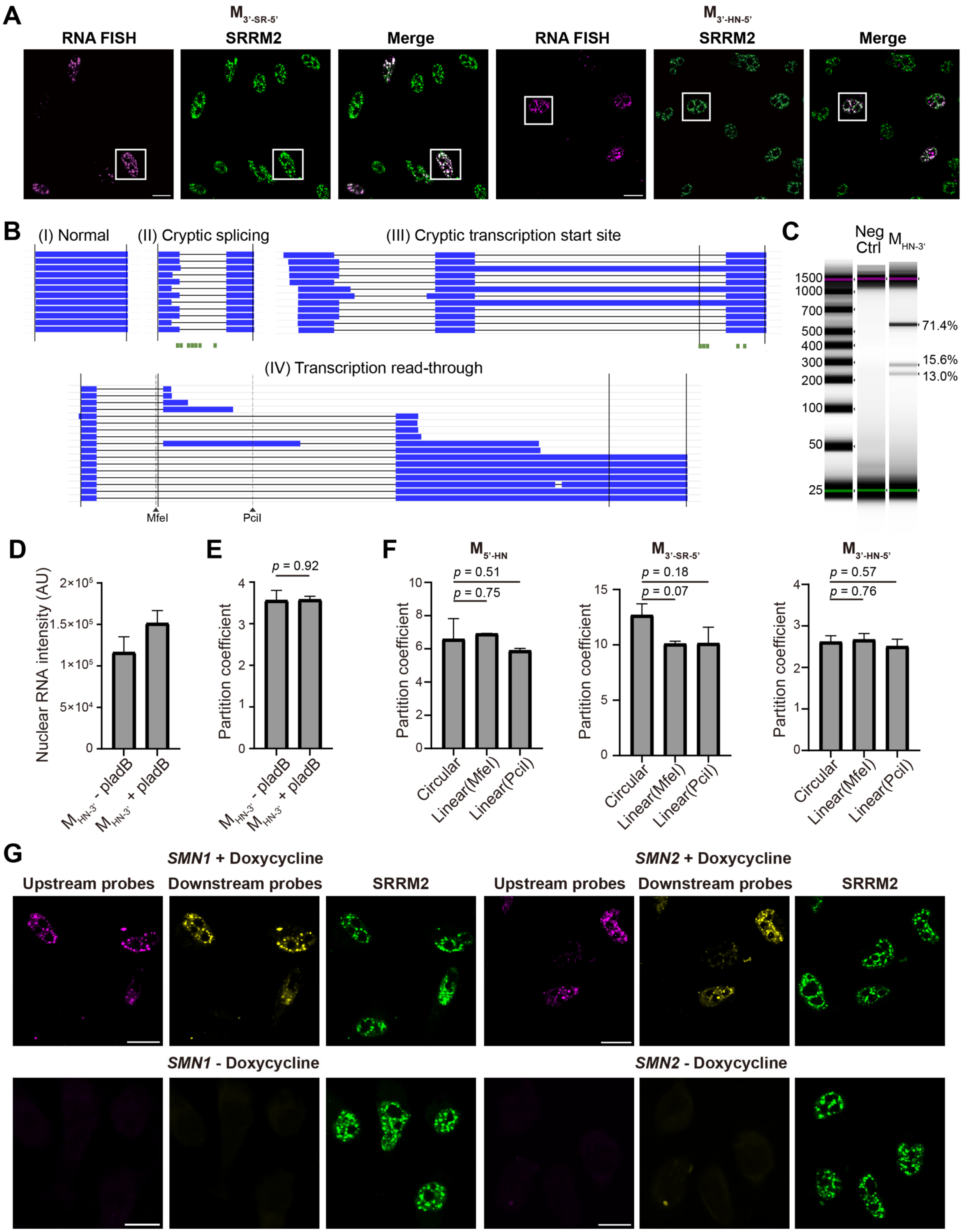


**Figure S2. Validations of constructs generating the unintended transcripts.**

(A) Representative zoom-out RNA FISH images of constructs $M_{3'\text{-SR-}5'}$ and $M_{3'\text{-HN-}5'}$ showing both transfected and untransfected cells. RNA FISH signals (labeled with AF647) are shown in magenta. Immunofluorescence signals of SRRM2 (labeled with AF488) are shown in green. Scale bar (white line) represents 20 μm. The boxed regions indicate the specific cells shown in Figure 1B-C.

(B) Illustration of the alignment of Nanopore sequencing reads demonstrating intended transcript (I), unwanted transcripts generated due to cryptic splicing (II), upstream cryptic transcription start site (III), and transcription read-through (IV). Green bars in (II) and (III) mark the FISH probe targeting positions. The intended transcript region is bounded by two thin vertical lines. Some transcripts in (IV) circle through the plasmid and include the intended transcription region again in their 3' end.

(C) Automated electrophoresis quantification of the fraction of unintended cryptic splicing product from $M_{HN-3'}$.

(D) Quantification of RNA FISH signal from $M_{HN-3'}$ construct in the cell nucleus in the absence and presence of Plad B treatment.

(E) Partition coefficient of $M_{HN-3'}$ in the absence and presence of Plad B treatment.

(F) Comparison of partition coefficients of $M_{5'-HN}$, $M_{3'-SR-5'}$, and $M_{3'-HN-5'}$ transcripts generated from circular plasmids and plasmids linearized using MfeI and PciI restriction enzymes.

(G) Representative RNA FISH images of *SMN1* and *SMN2* minimal constructs with and without doxycycline induction. RNA FISH signals using upstream probes (labeled with AF647) and downstream probes (labeled with CF568) are shown in magenta and yellow respectively. Immunofluorescence signals of SRRM2 (labeled with AF488) are shown in green. Scale bar (white line) represents 20 μm.

Error bars report the standard deviation from 2 biological replicates, with each replicate including 28–393 cells. *P*-values were calculated from unpaired two-tailed t-test.

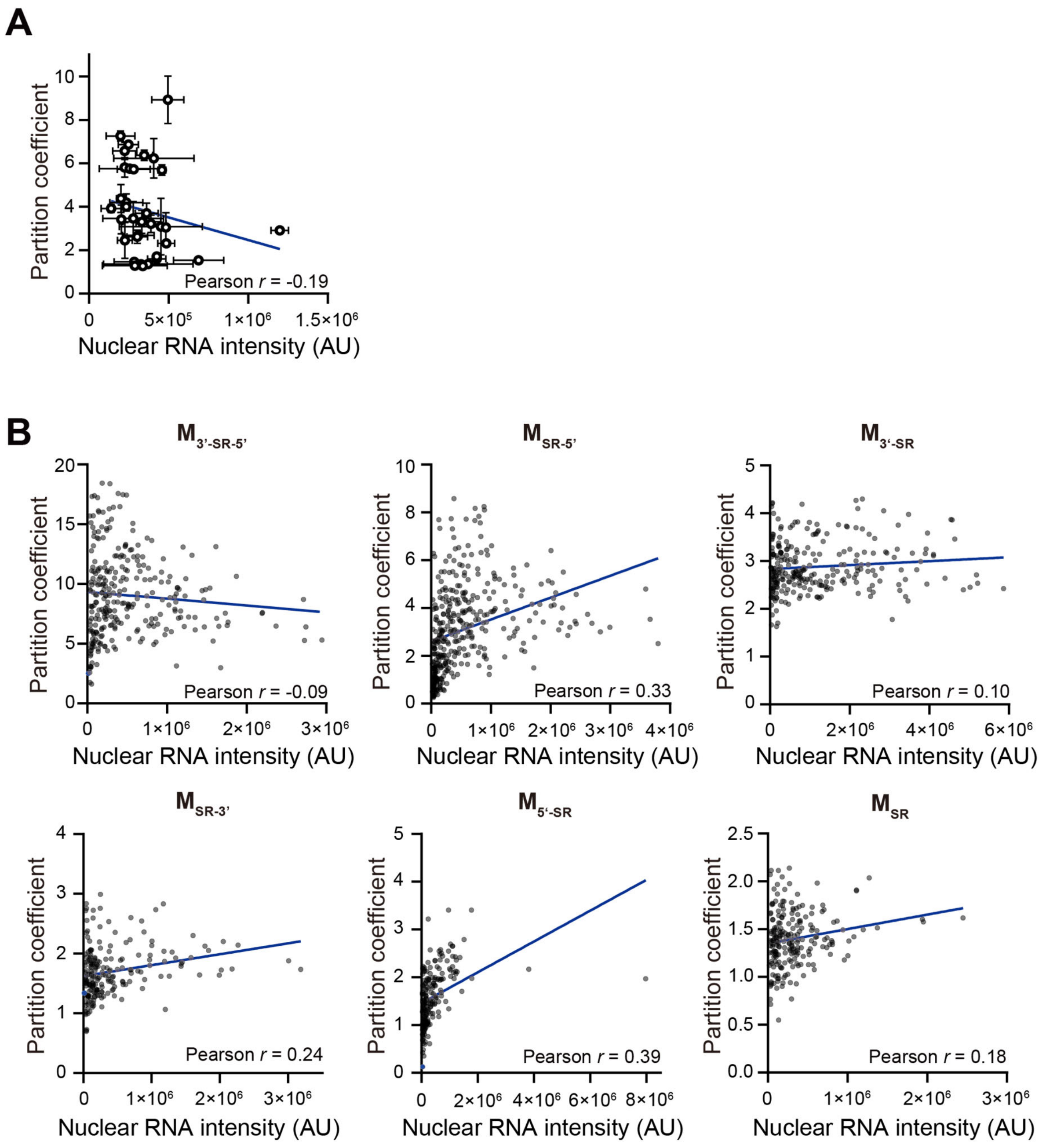


**Figure S3: Nuclear speckle localization is not driven by the nuclear abundance of RNA.**

(A) Correlation between population-averaged partition coefficient and nuclear abundance of RNA for all minimal constructs. Error bars report the standard deviation from 2-3 biological replicates, with each replicate including 34–840 cells.

(B) Correlation between partition coefficient and nuclear abundance of RNA at the single-cell level for single-module minimal constructs containing SR motifs. Each plot contains 192-375 cells from 2 biological replicates.

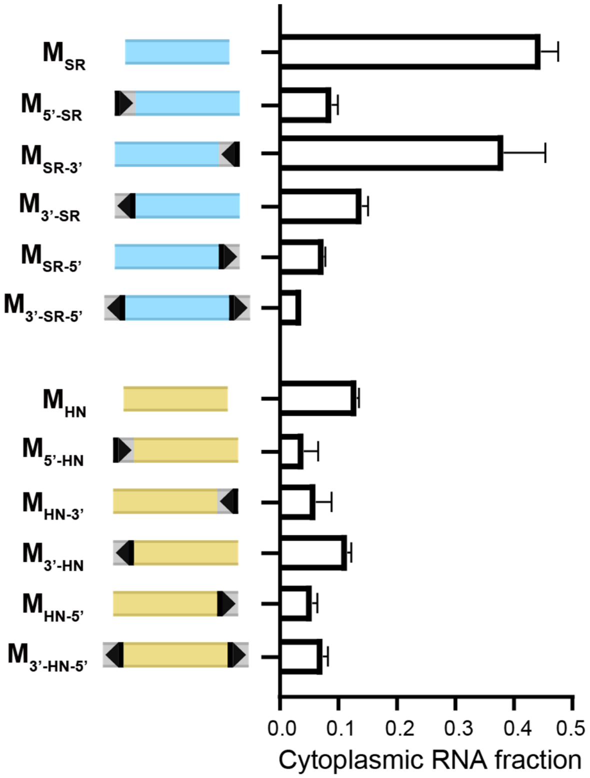


**Figure S4. Cytoplasmic fraction of minimal constructs.**
Fraction of RNA signal in the cytoplasm over total RNA signal in the cell for single-module minimal constructs. Error bars report the standard deviation from 2 biological replicates, with each replicate including 69–205 cells.

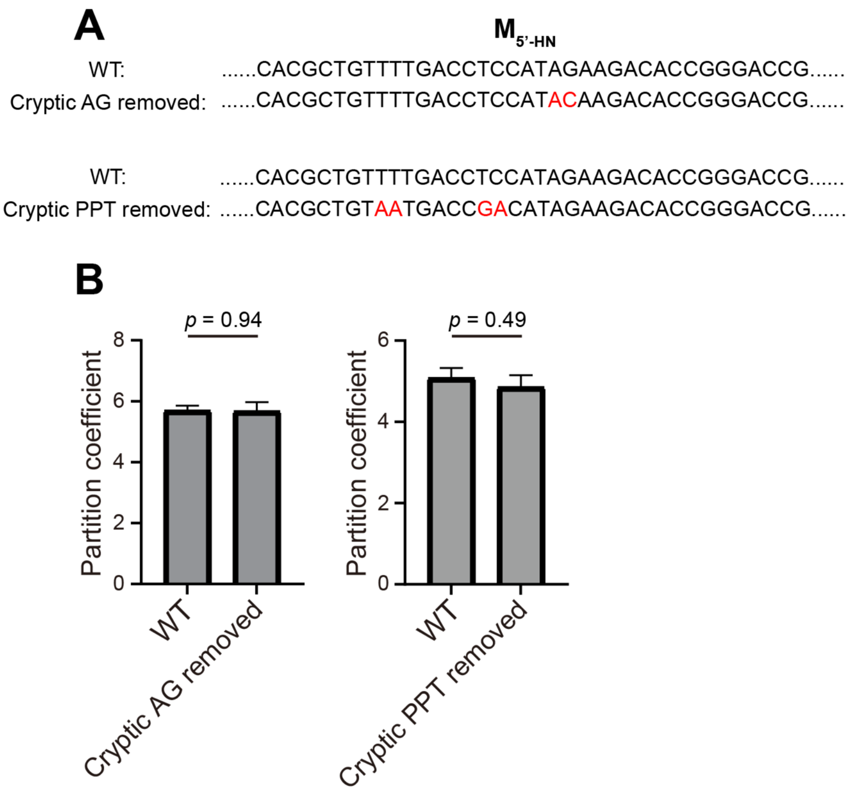


**Figure S5: Additional validation on $M_{5'\text{-HN}}$.**

(A) Schematic illustration of mutations introduced to $M_{5'\text{-HN}}$ to remove a 3' splice site-like AG sequence (cryptic AG removed) and a poly-pyrimidine tract-like sequence (cryptic PPT removed).

(B) Comparison of partition coefficients between WT and two mutant $M_{5'\text{-HN}}$. Error bars report the standard deviation from 2 biological replicates, with each replicate including 137–321 cells. *P*-values were calculated using unpaired two-tailed t-test.

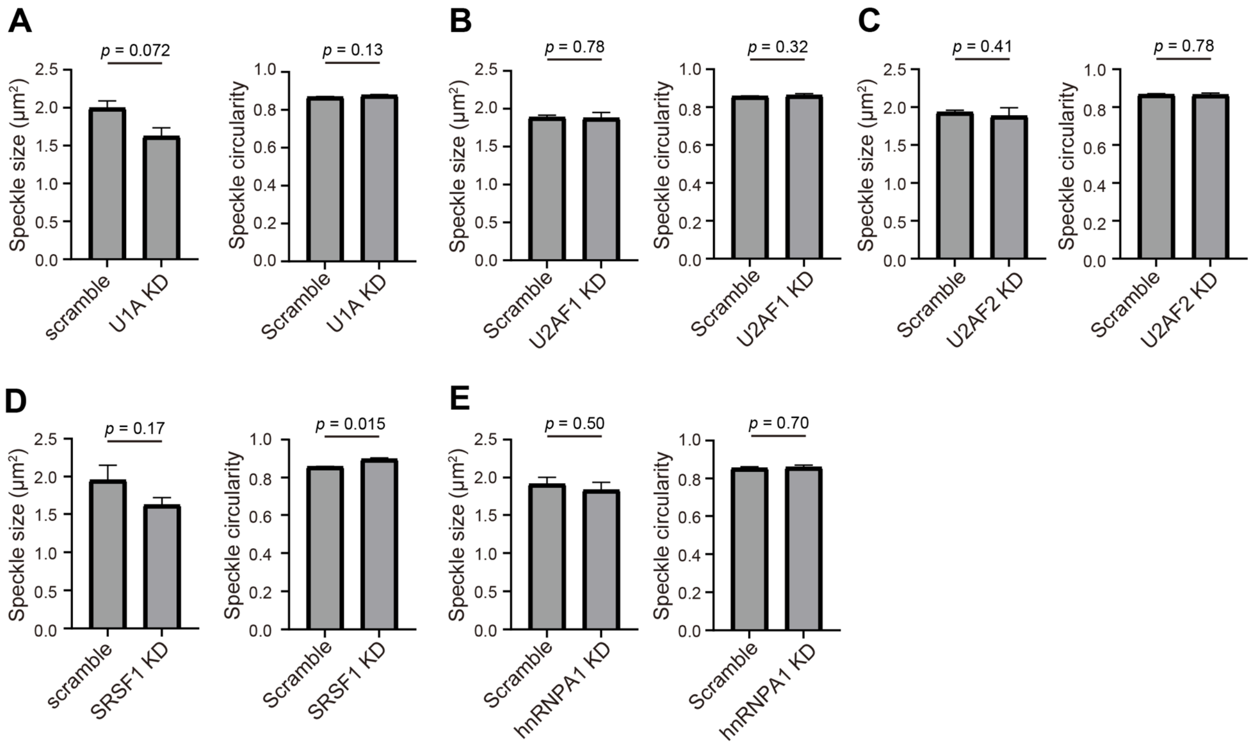


**Figure S6. Nuclear speckle morphology changes under all knockdown conditions.**

Changes in the speckle size and the circularity upon knockdown of (A) U1A, (B) U2AF1, (C) U2AF2, (D) SRSF1, and (E) hnRNPA1. The circularity was calculated as $\frac{4\pi A}{L^2}\left(1-\frac{0.5}{r}\right)^2$ where $r = \frac{L}{2\pi} + 0.5$, $A$ is the area, and $L$ is the perimeter. This yields a value between 0 and 1, with values closer to 1 indicating shapes that are more circular. Error bars report the standard deviation from 2 biological replicates, with each replicate including 137–321 cells. *P*-values were calculated from unpaired two-tailed t-test.

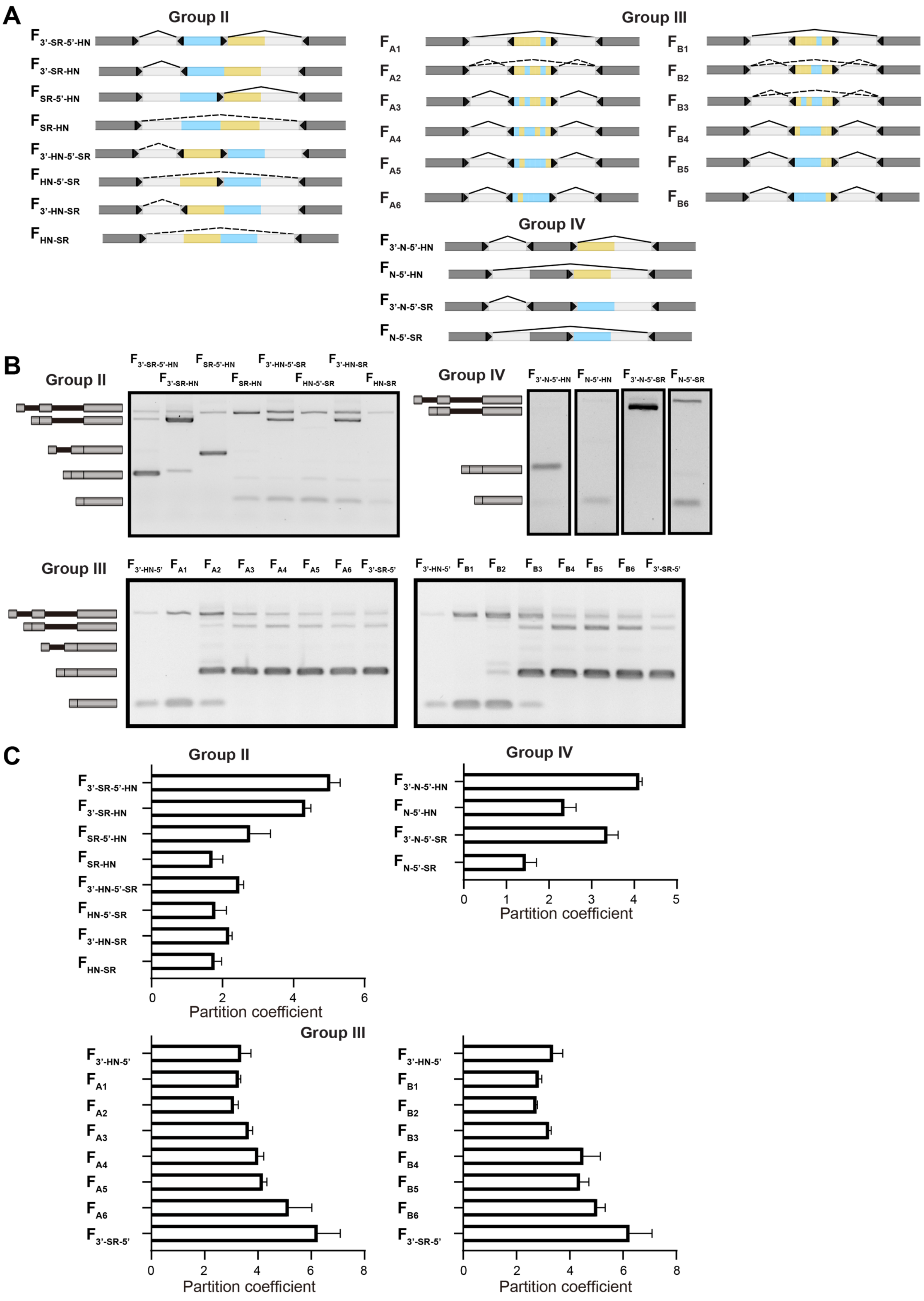

A
Group II
$F_{3'-SR-5'-HN}$
$F_{3'-SR-HN}$
$F_{SR-5'-HN}$
$F_{SR-HN}$
$F_{3'-HN-5'-SR}$
$F_{HN-5'-SR}$
$F_{3'-HN-SR}$
$F_{HN-SR}$
Group III
$F_{A1}$ $F_{A2}$ $F_{A3}$ $F_{A4}$ $F_{A5}$ $F_{A6}$
$F_{B1}$ $F_{B2}$ $F_{B3}$ $F_{B4}$ $F_{B5}$ $F_{B6}$
Group IV
$F_{3'-N-5'-HN}$
$F_{N-5'-HN}$
$F_{3'-N-5'-SR}$
$F_{N-5'-SR}$
B
Group II
Group IV
Group III
$F_{3'-HN-5'}$
$F_{3'-SR-5'}$
C
Group II
Group IV
Group III
Partition coefficient

**Figure S7. Analysis of additional minigene constructs.**

(A) Schematic illustration of minigene constructs containing dual-module minimal construct sequences (II), trajectory minimal construct sequences (III), and four additional minigene constructs containing neutral sequences (IV). Splicing outcomes are marked using solid lines (presenting major splicing products) or dashed lines (representing partial splicing products).

(B) RT-PCR gel electrophoresis analysis of splicing products for minigene constructs in (A).

(C) Partition coefficients of pre-mRNAs from minigene constructs in (A). Results for $F_{3'-SR-5'}$ and $F_{3'-HN-5'}$ are duplicated between both bar plots for an easy comparison. Error bars in the plots report the standard deviation (n=2) from two replicates, each replicate containing at least 60-90 nuclear speckles collected from 4-6 cells.
